# Single-Domain Antibody-Based Autophagosome-Targeting Chimera for Tau Clearance and Motor Function Restoration in Tauopathies

**DOI:** 10.1101/2025.06.30.662446

**Authors:** Yixiang Jiang, Amber M. Tetlow, Yan Lin, Changyi Ji, Klaudia F. Laborc, Adam C. Mar, Ruimin Pan, Xiang-Peng Kong, Erin E. Congdon, Einar M. Sigurdsson

## Abstract

Tauopathies are neurodegenerative diseases characterized by pathological tau accumulation, leading to motor and neuropsychiatric symptoms. Effective tau-targeting therapies remain a major challenge. Here, we present 1D9-LIRΔTP53INP2, a single-domain antibody (sdAb)-based protein degrader that facilitates tau clearance via the autophagy-lysosomal pathway. This engineered molecule combines the anti-tau sdAb 1D9 with an LC3-interacting region (LIRΔTP53INP2) to promote autophagosomal recruitment, mimicking autophagy receptors by simultaneously binding tau and LC3. In frontotemporal dementia (FTD) patient-derived neurons and JNPL3 tauopathy mice, both harboring the P301L tau mutation, 1D9-LIRΔTP53INP2 significantly reduced tau levels and improved motor function in mice. These findings underscore the therapeutic potential of sdAb-based protein degraders for tauopathies. Given the challenges of brain delivery for conventional antibodies, sdAbs with enhanced brain penetration and efficacy offer a promising strategy for treatment of neurodegenerative diseases.

## Background

Tauopathies, including Alzheimer’s disease (AD), progressive supranuclear palsy (PSP), corticobasal degeneration (CBD), and frontotemporal dementia (FTD), constitute a group of neurodegenerative diseases characterized by pathological tau protein accumulation in neurons and glia. Hyperphosphorylation of tau leads to its detachment from microtubules, promoting misfolding and aggregation into neurofibrillary tangles. Clinically, tauopathies manifest as cognitive and behavioral impairments, movement disorders, language deficits, and progressive memory decline. Despite extensive research, no effective treatments exist, highlighting the urgent need for advances in elucidating pathogenesis, biomarker discovery, and disease-modifying therapies ^1, 2^.

The tau protein is pivotal in tauopathies’ pathogenesis, with accumulating evidence indicating that post-translational modifications drive its mis-localization, oligomerization, and toxicity. While the precise molecular pathways through which tau induces neuronal toxicity and death remain incompletely elucidated, the clearance of toxic tau presents a promising therapeutic avenue to mitigate neuronal degeneration ^1, 3^.

Developing treatments for tauopathies by targeting the tau protein poses significant challenges, particularly due to the lack of a well-defined tau fold and active sites, as well as the intracellular localization of tau within the brain. Various strategies are under investigation ^1^, including: (1) active and passive immunization to facilitate tau clearance; (2) antisense oligonucleotides (ASO) to reduce tau synthesis; (3) enhancement of autophagy and proteasomal degradation pathways by using modulators; and (4) small-molecule inhibitors aimed at modulating tau aggregation. However, each approach is associated with potential limitations. For instance, immunization strategies may trigger immunological side effects ^4^, ASOs could lead to unintended gene silencing and delivery challenges ^5^, enhancing lysosomal or proteasomal activity may be hindered by pre-existing impairments and lack of specificity ^6, 7^, and small molecule inhibitors targeting protein aggregation may impact other proteins as well ^8^.

Tau pathology spreads throughout the brain via anatomically connected neurons. This may be caused by release and uptake of pathological tau via connected neurons but may also relate to tau pathology-induced alterations in neuronal activity ^9–11^. If the former, targeting extracellular tau for clearance may be beneficial. However, most of pathological tau is found inside neurons so it should ideally be targeted there ^12^. Although full-length IgG antibodies (∼150 kDa) can get into the brain and into neurons, their uptake and distribution within the brain and neurons is likely less than that of smaller antibody fragments.

To enhance efficacy, employing smaller antibody fragments like single-chain variable fragments (scFv, ∼25 kDa) or single-domain antibodies (sdAbs or VHHs, ∼15 kDa) is a promising approach. The reduced size of these fragments may facilitate improved blood-brain barrier (BBB) penetration and the targeting of cryptic epitopes inaccessible to full antibodies. Moreover, sdAbs exhibit enhanced stability and simplified large-scale production compared to whole antibodies. Notably, preclinical investigations involving scFvs and sdAbs in cellular and animal models have demonstrated their potential to impede tau polymer formation, serve as imaging agents, and mitigate tau pathology ^13–21^.

Reliable markers for assessing tau proteopathic burden and determining the stage of pathology in patients represent a significant challenge in the development of therapeutic interventions. In response to this challenge, we have recently introduced an sdAb-based in vivo imaging probe that enables specific and non-invasive visualization of tau pathology in mice, with the brain signal showing a strong correlation with lesion burden ^15^. By leveraging phage display libraries generated from a llama immunized with recombinant human tau (2N4R) and enriched paired helical filament (PHF) tau extracted from human tauopathy brain tissue, we identified two promising sdAb candidates, namely 2B8 and 1D9 ^17^. These sdAbs exhibit robust interactions with pathological tau protein from both human and mouse brain samples, underscoring their potential as diagnostic tools. Furthermore, apart from their application in imaging, we have explored the therapeutic potential of these sdAbs and investigated the impact of their four different IgG subclasses ^17^.

We and others have demonstrated that antibodies exhibit the ability to target proteinopathies both intracellularly and extracellularly ^17, 21–37^. Antibodies capable of penetrating neurons can specifically bind to protein aggregates within the endosomal-lysosomal system and/or ubiquitin-proteasome system, facilitating their clearance. Given tau’s predominantly intracellular localization, the intracellular targeting of this protein should be more efficacious than solely focusing on extracellular clearance. Nonetheless, the efficacy of the lysosome and proteasome, the two major intracellular protein degradation compartments, may be compromised in neurodegeneration and aging ^6, 7^. Dysfunction of these organelles could potentially exacerbate tau toxicity by hastening its accumulation owing to impaired protein degradation mechanisms.

An emerging strategy in drug development, PROteolysis TArgeting Chimeras (PROTACs) technology, has significantly broadened the scope of druggable proteins. PROTACs have been effectively employed to enhance tau clearance by upregulating the proteasomal pathway ^38–41^. Typically, small molecules or peptides serve as the ligands for the target, although they may exhibit low binding affinity and specificity and encounter challenges crossing the blood-brain barrier (BBB). Recently, two groups demonstrated the successful clearance of tau tangles in tauopathy mouse models by tagging the catalytic RING domain of the ubiquitin ligase TRIM21 to either anti-tau single-domain antibodies (sdAbs) or aggregation-prone tau. This approach leveraged the ubiquitin-proteasome system via adeno-associated virus (AAV) delivery of the protein degrader ^42, 43^. Despite these promising results, a key limitation remains: the proteasome’s narrow entry channel, which is primarily suited for degrading small, soluble proteins. As a result, aggregated proteins in higher-order structures cannot be cleared via this pathway, and these aggregates can obstruct proteasomal entry and impair the degradation of other proteins ^44^.

The autophagy-lysosomal pathway (ALP) serves as a specialized system for the degradation of large intracellular components, contrasting with the proteasome system ^44^. Ubiquitinated proteins are recognized by autophagy receptors through their ubiquitin-binding domains (UBD), and autophagy begins with the formation of double-membrane autophagosomes. Interaction between autophagy receptors and LC3 via the LC3-interacting region (LIR) facilitates phagophore formation and autophagosome closure, which subsequently fuse with lysosomes to degrade the enclosed substrates ^45^ (**Figure 1A**). Autophagy deficiencies, such as reduced LC3 expression and mutations in the ubiquitin-binding domain (UBD), have been implicated in various human diseases, particularly neurodegenerative disorders. These deficiencies impair the clearance of cytotoxic aggregates, disrupting cellular homeostasis ^46^. Consequently, accelerating or restoring ALP functions presents promising avenues for therapeutic intervention.

**Figure 1:**
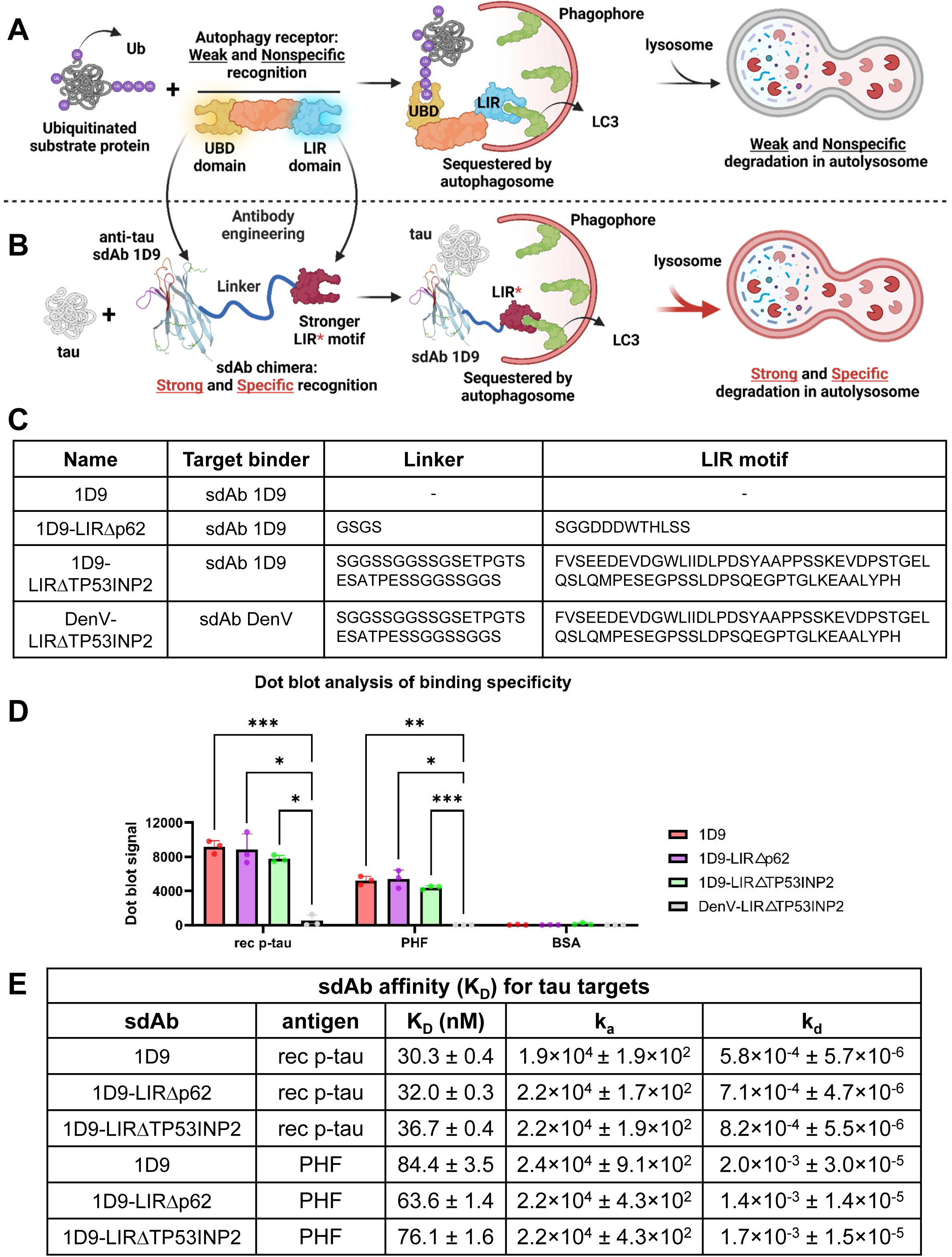
Design and characterization of single-domain antibody (sdAb)-based protein degraders for treatment of tauopathies. **(A)** Native autophagy process: Ubiquitinated proteins are recognized by autophagy receptors via their ubiquitin-binding domain (UBD). Interaction with LC3 through the LC3-interacting region (LIR) facilitates phagophore formation, autophagosome closure, and lysosomal degradation. **(B)** sdAb-AUTOTAC engineering: The UBD is replaced with an anti-tau sdAb (1D9) that specifically binds tau. Two distinct LIR motifs, linked by peptide linkers, drive tau targeting and degradation via the autophagy-lysosome pathway. **(C)** Structural and sequence details: Diagrams and amino acid sequences of the four sdAb constructs used in the study. **(D)** Dot blot analysis of binding specificity: Signals quantified for recombinant phosphorylated tau (rec p-tau), patient-derived paired helical filament (PHF) enriched tau, and bovine serum albumin (BSA). The original blot is shown in **Supplemental Figure 1A**. *, ***, ****: *p* < 0.05, 0.001, 0.0001 (Two-way ANOVA, Tukey’s post-hoc test). **(E)** Binding kinetics and affinity: Biolayer interferometry was performed using sdAb 1D9 and its chimeras loaded onto Ni-NTA biosensors. Binding affinities (K_D_), association (k_a_) and dissociation (k_d_) rates were determined using increasing concentration of the different tau preparations, including rec p-tau and PHF tau. The binding curves are shown in **Supplemental Figure 1B-G**.

Recent advancements in targeted degradation techniques via the ALP have shown promise ^47–49^, with successful applications observed in targeting misfolded tau both in cellular models and tauopathy mouse models with small molecules as tau binders ^50^. However, antibody-based tau binders are more attractive as therapy because of greater specificity for its target over small molecules.

During autophagy, autophagy receptors play a crucial role in facilitating the selection of autophagosomal substrates by simultaneously interacting with both the substrates and autophagosomal membranes ^51^. Notably, sequestosome-1 (SQSTM1/p62) stands out as a prominent mammalian autophagy receptor ^52^. The ubiquitin-associated (UBA) domain of p62 engages with ubiquitinylated (Ub) substrates, leading to the formation of aggregates specifically during selective autophagy. Subsequently, these aggregates are directed into the developing autophagosome through the interaction between the LC3-interacting region (LIR) of p62 and the LC3/GABARAP proteins on the autophagosomal membranes.

Capitalizing on the protein-protein interactions of p62 in the context of autophagy, we have devised a strategy to amplify therapeutic efficacy by integrating intracellular targeted degradation approaches within the autophagy-lysosome pathway. We present the development of a single-domain antibody-based autophagosome-targeting chimera (sdAb-AUTOTAC) tailored for tauopathies (**Figure 1B**). The design of sdAb-AUTOTAC stems from the protein engineering of sdAb-fusion degraders, culminating in a highly efficient mechanism to transport the tau protein into the autophagic pathway for subsequent degradation.

Our investigations demonstrate that an sdAb-AUTOTAC protein degrader, named 1D9-LIRΔTP53INP2, enhances tau clearance in both patient-induced pluripotent stem cells (iPSC)-derived neurons and mouse tauopathy models by facilitating its degradation within the autolysosomes. These results underscore the potential of our sdAb-AUTOTAC protein degrader as a promising therapeutic candidate for tauopathies. Notably, considering the limited brain penetration of conventional antibodies, the development of more potent sdAbs capable of enhanced brain permeability could further amplify the clinical advantages of antibody-based therapeutic candidate.

## Materials and Methods

### Mammalian expression and purification of single domain antibodies and their engineered derivatives

The single domain antibodies (sdAbs) and their engineered derivative clones, anti-tau 1D9, 1D9-LIRΔp62, 1D9-LIRΔTP53INP2, anti-dengue virus (DenV) sdAb, and DenV-LIRΔTP53INP2, were expressed in a mammalian system to avoid having to endotoxin purify clones expressed in a bacterial system. pVRC8400-sdAb constructs were made by inserting their individual gene sequences between the 5’ EcoRI and 3’ AfeI sites with a signal peptide, mouse interleukin-2 (IL-2) leader sequence (MYRMQLLSCIALSLALVT) at N-terminus and 6xhistidine (His) tag at C-terminus for detection and purification. The sdAbs were then expressed in FreestyleTM 293F cells (Invitrogen, Cat. No. R790-07). Briefly, FreestyleTM 293F cells were transiently transfected with the mixture of DNA plasmid and polycation polyethylenimine (PEI; 25 kDa linear PEI, Polysciences, Inc., cat. No. 23966). The transfected cells were incubated at 125 rpm in FreestyleTM 293 Expression Medium for suspension culture at 37°C with 5% CO_2_. The supernatants were harvested 5 days after transfection, filtered through a 0.45 μm filter, followed by sdAb purification using Ni-NTA columns (GE Healthcare). The wash buffer consisted of 20 mM Na_3_PO_4_, 100 mM NaCl, and 20 mM imidazole, pH 7.4. The elution buffer consisted of 20 mM Na_3_PO_4_, 100 mM NaCl, and 500 mM imidazole, pH 7.4. Following elusion, the sdAb was dialyzed into PBS and its concentration determined by a bicinchoninic acid (BCA) assay.

### Preparation of recombinant hyperphosphorylated tau for binding assay

Recombinant phosphorylated tau (rec p-tau) was purified as previously described ^15, 17, 53^. Briefly, 1N4R tau isoform was co-expressed with GSK-3β in E. coli using the PIMAX approach. After induction with IPTG, cells were lysed, and the lysate was heat-treated to enrich p-tau. The supernatant was digested with TEV protease, followed by centrifugation and concentration. The sample was purified by size exclusion chromatography, pooled, concentrated, and stored in 10% glycerol at −80°C.

### Isolation of paired helical filament tau for binding assay

Paired helical filament (PHF)-enriched tau was extracted from human brain tissue samples collected from patients with mixed AD/Progressive supranuclear palsy (PSP) pathology (Case ID: PSP-65484-01-slab 3), sourced from the National Disease Research Interchange in Philadelphia, PA using methods described previously ^22, 32^. The selection of these samples was based on their proven ability to induce significant toxicity in preliminary studies involving primary neuronal and mixed neuron/glia cultures. The brain tissue was initially homogenized in a buffer solution containing 0.75 M NaCl, 1 mM EGTA, 0.5 mM MgSO_4_, and 100 mM 2-(N-morpholino) ethanesulfonic acid at pH 6.5, followed by centrifugation at 11,000 x g for 20 minutes. The resulting low-speed supernatant was treated with 1% sarkosyl (prepared from a 10% stock solution in PBS) for one hour at room temperature (RT). Subsequently, the mixture underwent centrifugation at 100,000 x g for 60 minutes, and the supernatant was carefully decanted. The pellet was washed with a 1% sarkosyl solution before the tau protein was resolubilized by briefly heating to 37°C in 50 mM Tris-HCl buffer, followed by overnight dialysis in PBS for purification.

### Dot blot assay

Rec p-tau, PHF-tau, and bovine serum albumin (BSA) (5 µL of 1 mg/mL each) were spotted onto a 0.2 µm nitrocellulose membrane and air-dried for 30 minutes. The membrane was then blocked for 1 hour at RT with 5% non-fat milk in TBS-T (0.1% Tween-20 in Tris-buffered saline).

Following blocking, the membrane was incubated overnight at 4 °C with sdAbs: 1D9, 1D9-LIRΔp62, 1D9-LIRΔTP53INP2, or DenV-LIRΔTP53INP2, at a concentration of 600 nM in SuperBlock™ buffer (Thermo Fisher Scientific). All sdAbs contained a 6xHis tag for detection.

After several washing with TBS-T, the membrane was incubated for 1 hour at RT with an anti-6xHis tag primary antibody (1:2000, ABCAM, clone EPR20547) diluted in SuperBlock™. This was followed by additional washes with TBS-T and incubation with an IRDye® 800CW-conjugated secondary antibody (1:10,000, LI-COR Biosciences).

Immunoreactive signals were visualized and quantified using the LI-COR Image Studio Lite software (version 5.2).

### Bio-layer interferometry binding affinity assay

The bio-layer interferometry (BLI) experiments were carried out using a ForteBio Octet® RED96 instrument in a 96-well black flat bottom plate with shaking speed set at 1,000 rpm, at RT. Samples were prepared in 200 μL of 1X PBS assay buffer containing 0.05% Tween-20 at pH 7.4.

To measure the biophysical interaction between the His-tagged sdAbs and the tau protein, we used the nickel (Ni)-NTA biosensor, which is pre-immobilized with Ni-charged tris-nitrilotriacetic (Tris-NTA) groups to capture His-tagged molecules. Before the BLI analysis, the sensor was hydrated in the assay buffer for 10 min, and a baseline was established in the buffer for 120 sec. The Ni-NTA tips were then loaded with His-tagged sdAbs at 300 nM for 120 sec, resulting in a specific interaction between the His-tag and Ni ions. Another baseline step was performed in the assay buffer for 120 sec, after which the ligand-loaded biosensor tips were dipped into the tau antigen solutions at different concentrations in 1X PBS (pH 7.4) with 0.05% Tween-20 (PBS-T) for 300 sec. Finally, dissociation was conducted in the assay buffer for 400 sec. A reference biosensor was loaded with the His-tagged sdAb and ran with an assay buffer blank for the association and dissociation steps.

Data analysis was performed using Data Analysis 11.0 software with reference subtraction using a 1:1 binding model as we have described previously in detail ^15^.

### Human neuronal cell culture and sdAb treatment

Induced pluripotent stem cell (iPSC)-derived human neurons used in this study were established from the Tau Consortium iPSC line collection ^54^, grown at NSCI core facility NeuraCell, and are available upon request (www.neuralsci.org/tau). iPSCs were generated from fibroblasts obtained from an individual carrying the autosomal dominant P301L mutation (Donor ID: F0510) (**Figure 2A**). Fibroblasts were reprogrammed into iPSCs and subsequently differentiated into cortical-enriched neural progenitor cells (NPCs) and neurons, as previously described ^54^.

**Figure 2:**
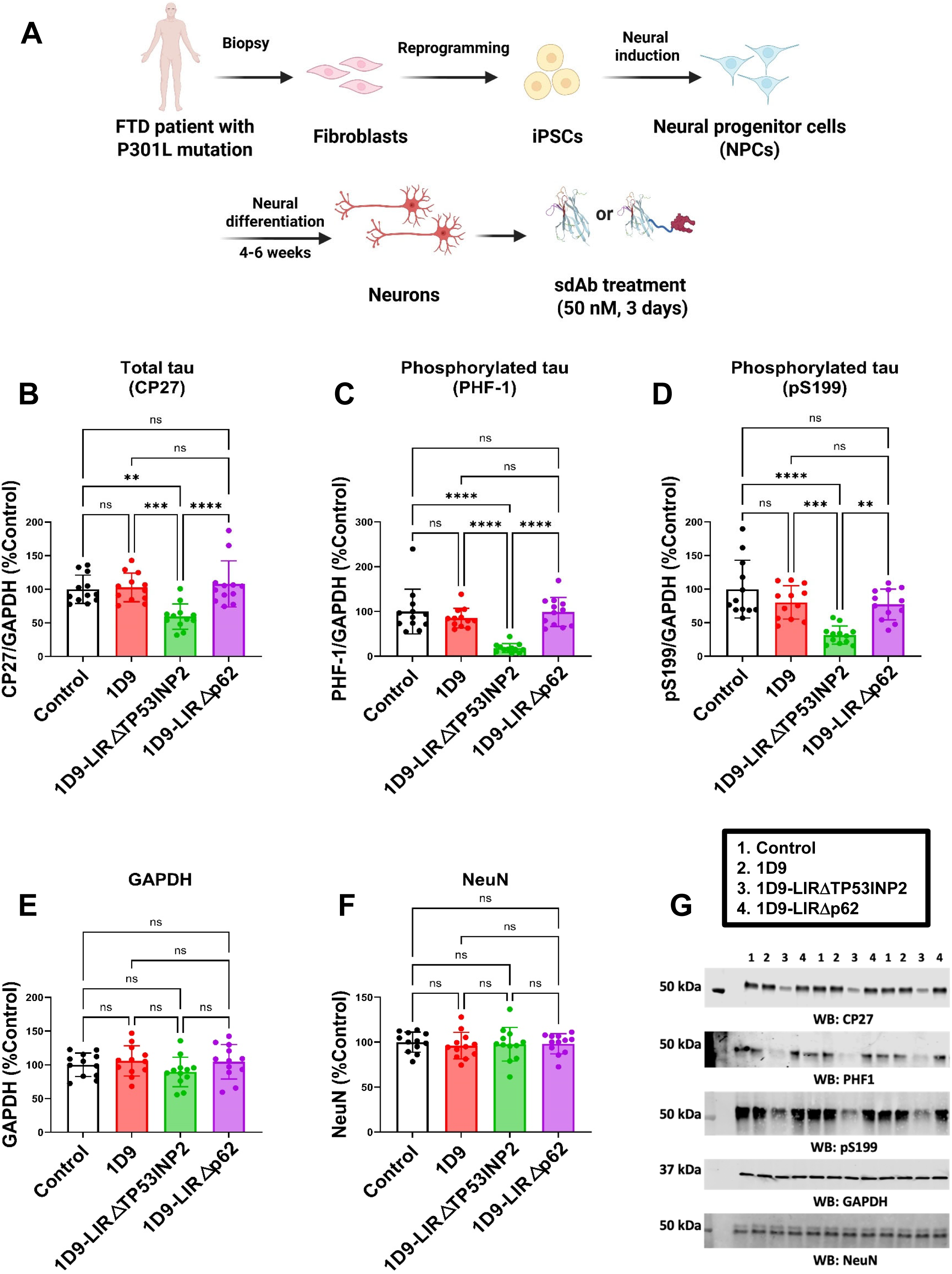
Enhanced intracellular clearance of tau by modified sdAb 1D9-LIRΔTP53INP2 chimera in a human neuronal cell model of tauopathy. **(A)** Diagram for generation of patients’ iPSC-derived cortical-enriched neural progenitor cells (NPCs) and differentiation into mature cortical neurons, followed by administration of sdAb treatment (50 nM) or vehicle control, with samples collected 72 h post-antibody application (*n* = at least 12 replicates per condition). **(B)** One-way ANOVA revealed significant differences in normalized total tau levels (CP27, p < 0.0001). 1D9-LIRΔTP53INP2 treatment showed 41% (*p* = 0.0011), 43% (*p* = 0.0005), and 49% (*p* < 0.0001) decrease compared to control, 1D9, and 1D9-LIRΔp62 treatment groups, respectively. **(C)** One-way ANOVA revealed significant differences in normalized p-tau levels (PHF1, p < 0.0001). 1D9-LIRΔTP53INP2 treatment showed 82% (*p* < 0.0001), 67% (*p* < 0.0001), and 81% (*p* < 0.0001) decrease compared to control, 1D9, and 1D9-LIRΔp62 treatment groups, respectively. **(D)** One-way ANOVA revealed significant differences in normalized pSer199 tau levels (p < 0.0001). 1D9-LIRΔTP53INP2 treatment showed 68% (*p* < 0.0001), 49% (*p* = 0.0006), and 46% (*p* < 0.0014) decrease compared to control, 1D9, and 1D9-LIRΔp62 treatment groups, respectively. **(E-F)** One-way ANOVA revealed no significant differences in GAPDH levels (*p* = 0.2706) or NeuN levels (*p* = 0.9283). **(G)** Representative immunoblots illustrating total tau (CP27), phosphorylated tau (PHF1 and pSer199), GAPDH and NeuN levels in treated human neuronal cell lysates. The complete western blots are shown in **Supplemental Figure 2**. **, ***, ****: p < 0.01, 0.001, 0.0001 (Tukey’s post-hoc test).

Tissue culture plates were coated with poly-L-ornithine (PLO) and laminin to enhance cell adhesion. PLO was diluted to 15 µg/mL in PBS, applied evenly to the culture surface, and incubated for at least 2 hours at RT or overnight at 2–8°C (sealed). Following incubation, excess PLO was removed, and wells were washed twice with PBS and once with DMEM/F-12 (Dulbecco’s Modified Eagle Medium/Nutrient Mixture F-12). Laminin was then diluted to 10 µg/mL in DMEM/F-12, added to the PLO-coated plates, and incubated under the same conditions. Coated plates were either used immediately or stored at 2–8°C (sealed) for up to 2 weeks and equilibrated to RT for 30 minutes before cell plating.

NPCs were maintained and expanded in STEMdiff™ Neural Progenitor Medium (Basal Medium [#05834] + Supplement A [#05836] + Supplement B [#05837]), following manufacturer’s protocol #28782 (version 2.4.0). NPCs were passaged at densities of 100,000 cells per well in 6-well plates, 50,000 cells per well in 12-well plates, or 25,000 cells per well in 24-well plates and used for differentiation before passage four to minimize astrocyte contamination and maintain neuronal differentiation efficiency. To induce terminal differentiation, NPCs (≤ passage 4) were plated in BrainPhys™ Neuronal Medium supplemented with the N2-A & SM1 Kit (#05793), 20 ng/mL GDNF, 20 ng/mL BDNF, 1 mM dibutyryl-cAMP, and 200 nM ascorbic acid. Cells were differentiated for 6–8 weeks with media changes performed twice weekly, following protocol #10000000225 (version 04).

To evaluate the efficacy of sdAb constructs in tau clearance, iPSC-derived human neurons were treated with sdAbs in 24-well plates. Treatment was conducted in a total volume of 0.5 mL per well by replacing 0.25 mL of conditioned media with an equal volume of fresh media pre-mixed with sdAbs at a 2× working concentration. Cells were incubated at 37°C for 3 days before analysis.

To investigate the degradation mechanism, lysosomal and proteasomal pathways were selectively inhibited using bafilomycin A1 (Baf.A1) and MG132, respectively. Neurons were pre-treated with either the lysosome inhibitor Baf.A1 (0.2 µM) or the proteasome inhibitor MG132 (2 µM) for 6 hours. Subsequently, the sdAb construct (25 nM) was added directly to the media without media exchange. Cells were incubated at 37°C for 2 days before analysis.

### Immunofluorescence assay

iPSC-derived human neurons were cultured on pre-coated coverslips as previously described. To inhibit lysosomal degradation, neurons were pre-treated with bafilomycin A1 (Baf.A1, 0.2 µM) for 6 hours. The sdAb construct (25 nM) was then added directly to the culture medium without media exchange, and cells were incubated at 37°C for 24 hours.

Cells were fixed with 4% (v/v) paraformaldehyde in HBSS (Hanks’ balanced salt solution) containing 4% sucrose for 20 minutes at RT, followed by three washes with HBSS (10 minutes each). Permeabilization and blocking were performed in HBSS containing 0.2% Triton X-100, 5% bovine serum albumin (BSA), 2% normal donkey serum (NDS), and 0.3 M glycine for 2 hours at RT. The cultures were then gently washed twice with HBSS (5 minutes each) to minimize cell detachment.

Cells were incubated overnight at 4°C with primary antibodies: anti-tau (AT8, MN1020, 1:100), anti-HIS tag (PA1-9531, 1:100), and anti-LC3 (PA1-46286, 1:100) in HBSS containing 5% BSA and 1% NDS. Following two washes with HBSS (5 minutes each), cells were incubated with secondary antibodies at RT for 1 hour: Alexa Fluor™ Plus 555 goat anti-chicken IgY (A32932, 1:1000), Alexa Fluor™ 647 goat anti-rabbit IgG (AB150083, 1:1000), and Alexa Fluor™ 488 goat anti-mouse IgG (AB150113, 1:1000) in HBSS containing 5% BSA and 1% NDS. Samples were then rinsed four times with HBSS. Finally, coverslips were mounted using ProLong™ Gold antifade reagent, and imaging was performed using a Zeiss LSM 800 confocal laser scanning microscope (Axioimager).

To quantify the co-localization of tau and LC3B, the Manders’ correlation coefficient was calculated using the JACoP plugin in ImageJ as previously described ^15^. The Manders’ coefficient measures the proportion of overlapping fluorescence signals, ranging from 0 (no co-localization) to 1 (complete co-localization).

### Co-immunoprecipitation

Neural progenitor cells (NPCs) were plated and differentiated for 6–8 weeks in pre-coated 6-well plates as previously described. For each co-immunoprecipitation (Co-IP) experiment, neurons from two wells were washed with PBS, harvested, and combined into a single pellet. To stabilize and detect protein complex formation, all steps were performed on ice or at 4°C.

Neurons were pre-treated with bafilomycin A1 (Baf.A1, 0.2 µM) for 6 hours, followed by the direct addition of the sdAb construct (25 nM) to the culture medium without media exchange. Cells were incubated at 37°C for 48 hours before analysis.

Cells were lysed in ice-cold Pierce™ IP Lysis Buffer (Thermo Fisher Scientific) for 15 minutes at 4°C, followed by centrifugation at 10,000 × g for 10 minutes. The resulting supernatant (Input) was collected, and protein concentration was determined using a BCA assay.

Co-IP was performed using the Thermo Scientific™ Pierce™ Crosslink Magnetic IP/Co-IP Kit (Cat. #88805) following the manufacturer’s protocol. Briefly, Protein A/G magnetic beads were pre-washed with 1× Coupling Buffer. Antibodies (10 µg of TAU5, LC3B, or control IgG) were diluted to a final volume of 100 µL and incubated with the beads for 15 minutes at RT. After three washes with 1× Coupling Buffer, the antibodies were cross-linked to the beads using disuccinimidyl suberate (DSS) for 30 minutes. The beads were then washed three times with Elution Buffer and twice with IP Lysis/Wash Buffer.

Lysates (200 µL, 300 µg total protein) were incubated with antibody-bound beads overnight at 4°C with gentle rotation. The beads were washed twice with IP Lysis/Wash Buffer and once with purified water. Bound proteins were eluted using 100 µL of Elution Buffer and incubated for 5 minutes at RT with rotation. Magnetic separation was performed, and the supernatant containing the eluted target antigen was collected. Samples were mixed with 4× Laemmli Sample Buffer (Bio-Rad), and 15 µL of each sample was loaded onto SDS-PAGE for western blot analysis.

Control western blot analysis with 2% (6 μg) of IP input confirms the treatment effect of sdAb + the lysosome inhibitor Baf.A1.

### Mouse models and sdAb treatment

Homozygous JNPL3 female mice, which express the human tau 0N4R isoform with the P301L mutation under the control of the mouse prion promoter, were used for in vivo studies. The transgene is expressed at approximately twice the level of endogenous mouse tau and is distributed throughout the brain, with the most severe tau pathology typically observed in the brainstem ^55^. However, extensive tau pathology is also present in other regions, including the cortex. Originally, homozygous JNPL3 mice exhibited severe motor deficits by approximately 10 months of age, likely due to tau pathology in the brainstem and spinal cord.

To assess baseline gait abnormalities in tauopathy mice, CatWalk™ XT automated gait analysis was conducted on a cohort of untreated 8-month-old female mice, including JNPL3 tauopathy mice (*n* = 9) and age-matched wild-type littermates (*n* = 7). This time point was selected to match the age of animals analyzed in the post-treatment cohort. Gait parameters were quantified using CatWalk™ software. These baseline measurements served as a reference for interpreting motor deficits in JNPL3 mice and for evaluating the efficacy of therapeutic interventions.

A total of 25 female JNPL3 mice, aged 6–7 months, were enrolled and randomly assigned to one of three treatment groups: vehicle control (*n* = 7), 1D9 (*n* = 9), or 1D9-LIRΔTP53INP2 (*n* = 9). Mice received six weekly intravenous (i.v.) injections of either vehicle control, unmodified 1D9 sdAb (200 nmol/kg, ∼3.2 mg/kg), or modified 1D9-LIRΔTP53INP2 sdAb (200 nmol/kg, ∼4.9 mg/kg) in PBS. Motor behavior was assessed following treatment using the CatWalk™ automated gait analysis system to determine the impact of each intervention on locomotor function.

### In Vivo Imaging System (IVIS) imaging and immunofluorescence staining

In vivo imaging was performed using the IVIS Lumina XR system (PerkinElmer) in homozygous P301L transgenic mice (female, 8 months old) as we have described previously in detail ^15, 21^. Before imaging, mice were anesthetized with 2% isoflurane and maintained with 1.5% isoflurane in 30% oxygen. VivoTag 680XL (PerkinElmer)-labeled sdAb constructs were administered via i.v. injection at a dosage of 700 nmol/kg (∼11.1 mg/kg for 1D9 and ∼16.9 mg/kg for 1D9-LIRΔTP53INP2). To minimize non-specific light diffraction, fur on the head and body was shaved prior to imaging.

Immediately after i.v. injection, fluorescence imaging was performed at defined intervals (every 15 minutes for 1 hour). Mice remained anesthetized within the imaging chamber, receiving a continuous supply of 2% isoflurane in oxygen (0.4 L/min) via a nose cone. The body temperature was maintained at 37°C using a thermoelectrically controlled imaging stage.

Fluorescent images were acquired using Living Image software (PerkinElmer) with the following settings: mode: fluorescent, illumination: epi-illumination, exposure time and binning: auto (30 s under these conditions), excitation filter: 675 nm, emission filter: Cy5.5 (695–770 nm). All acquired images were automatically time- and date-stamped for analysis.

Images were analyzed using Living Image software (PerkinElmer). Image sequences from different time points were loaded together, and the same color scale was applied to all images for consistent visual comparison. A circular region of interest (ROI) was defined over the brain region of each mouse. The calibrated radiant efficiency (photons/sec/cm²/sr/mW/cm²) was recorded for each ROI. Data was compiled into a table and used to generate IVIS imaging profiles.

After imaging, mice were transcardially perfused with PBS. The brains were dissected, and the right hemisphere (with a small portion of the left hemisphere to maintain structural integrity) was immersion-fixed overnight in 2% paraformaldehyde/lysine/periodate (PLP) at 4°C. The left hemisphere was snap-frozen on CO₂ blocks and stored at −80°C for subsequent western blot analysis.

Following overnight fixation, the right hemisphere was transferred to 2% dimethyl sulfoxide (DMSO) in 20% glycerol phosphate buffer and stored at 4°C until sectioning. Coronal brain sections (40 μm) were obtained and processed for immunofluorescence to detect 680XL-sdAb signals and assess subcellular localization via co-staining with phosphorylated tau (PHF1 antibody) and the autophagosome marker LC3B.

Immunofluorescence staining was performed on free-floating sections using a standard protocol. Briefly, sections were washed in PBS, permeabilized with 0.3% Triton X-100, and blocked in 5% BSA, 2% normal donkey serum (NDS), and 0.3 M glycine for 2 hour at RT. Sections were then incubated overnight at 4°C with primary antibodies against p-tau (PHF1) and LC3B (autophagosome) at a 1:100 dilution. Bound antibodies were detected using Alexa Fluor 488-conjugated goat anti-mouse/rabbit IgG (Invitrogen). After mounting with ProLong Gold antifade reagent, sections were imaged using a Zeiss LSM 800 confocal laser scanning microscope (Axioimager).

### Gait analysis

Gait was assessed through quantitative analysis of footfalls and parameters of quadrupedal locomotion in freely moving animals using the Catwalk XT (software version 10.7.704, Noldus Information Technology). For each trial, individual mice were placed on one end of the enclosed runway (200-cm length, 10-cm width, 20-cm height) and allowed to traverse unrestrained at their own pace. Mice explored the runway until they achieved a minimum of three compliant runs; runs in which they crossed the middle 60 cm of the runway within a 0.5-s to 10-s duration, at a minimum velocity of 5 cm/s and having speed variation less than 60%. A high-speed camera located underneath the transversely illuminated glass runway captured detailed data for each footfall. All footfalls were validated manually after automated classification. Detailed gait parameters for each mouse were then automatically computed and extracted based on the digital paw prints acquired during the runs. Body weight and average run speed were included as covariates in the statistical analysis whenever they exerted a significant impact on the outcome measure of interest. All statistical analyses were performed using IBM SPSS Statistics (version 28.0.0.1).

### Brain homogenization, tau fraction preparation, and western blot analysis

Following the behavioral experiment, mice were transcardially perfused with PBS. The brains were collected for biochemical and immunohistochemical analysis. As described previously ^22^, the left hemisphere of each brain was homogenized in modified RIPA buffer (50 mM Tris-HCl, 150 mM NaCl, 1 mM EDTA, 1% Nonidet P-40, pH 7.4) supplemented with protease and phosphatase inhibitors, including a 1X protease inhibitor mixture (cOmplete, Roche), 1 mM sodium fluoride (NaF), 1 mM sodium orthovanadate (Na₃VO₄), 1 nM phenylmethylsulfonyl fluoride (PMSF), and 0.25% sodium deoxycholate. Homogenates were briefly kept on ice and centrifuged at 20,000 × g for 20 minutes at 20°C. The resulting supernatant, designated as the low-speed supernatant (LSS), was collected as the soluble tau fraction. Protein concentrations were determined using the BCA assay, and samples were adjusted to equal concentrations with homogenization buffer. A total of 5 μg of protein was loaded per lane for further analysis.

To obtain the sarkosyl-insoluble tau fraction, equal amounts of protein (2 mg) from the LSS were mixed with 10% sarkosyl in phosphate buffer to reach a final concentration of 1% sarkosyl. The mixture was incubated for 30 minutes at RT on a Mini LabRoller (Labnet H5500), followed by ultracentrifugation at 100,000 × g for 1 hour at 20°C using a Beckman Coulter Optima™ MAX-XP Ultracentrifuge. The resulting pellet (sarkosyl pellet (SP), sarkosyl insoluble fraction) was washed with 1% sarkosyl solution and centrifuged again at 100,000 × g for 1 hour at 20°C. The final pellet was air-dried for 30 minutes, then resuspended in 50 μL of modified O+ buffer containing 62.5 mM Tris-HCl, 10% glycerol, 5% β-mercaptoethanol, 2.3% SDS, 1 mM EDTA, 1 mM EGTA, 1 mM NaF, 1 mM Na₃VO₄, 1 nM PMSF, and a 1X protease inhibitor mixture. Samples were boiled before loading 10 μL per lane on 12% SDS-PAGE for electrophoresis.

Following protein transfer to nitrocellulose membranes, blots were blocked with SuperBlock® (Thermo Fisher Scientific) and incubated overnight at 4°C with primary antibodies, including CP27 (1:500, gift from P. Davies) for total tau detection, PHF1 (1:1000, gift from P. Davies), pSer199 (1:1000, Invitrogen 44-734G) and AT8 (1:1000, MN1020) for p-tau detection, GFAP (1:1000, PA5-16291), Iba-1 (1:1000, 10904-1-AP) and GAPDH (1:4000, Cell Signaling Technology, D16H11) as a loading control. After incubation, membranes were treated with IRDye® 800CW or IRDye® 680RD secondary antibodies (1:10,000, LI-COR Biosciences). Immunoreactive bands were visualized and quantified using LI-COR Image Studio Lite 5.2 software.

### Immunohistochemistry

As described previously in detail ^56^, the fixed right-brain hemisphere was cut coronally into 40 µm thick sections, using a freezing cryostat, and stored at −20°C in ethylene glycol cryoprotectant until retrieved for immunohistochemistry. Prior to staining, free-floating sections were washed in PBS to remove cryoprotectant.

### PHF1 and MC1 staining

Tissue sections were placed in a free-floating staining rack with 0.3% hydrogen peroxide in Tris buffered saline (TBS, pH 7.4) with 0.25% Tween 20 to block endogenous peroxidases and to permeabilize the sections. Non-specific binding sites were then blocked for 1 hour at RT using 5% powdered milk in TBS. After TBS-T (0.05% Tween-20) washes, sections were incubated overnight at 4°C with primary antibodies diluted in TBS: PHF1 (1:2500) and MC1 (1:100) (both gifts from Peter Davies). The following day, tissues were washed with TBS containing 0.05% Tween 20. The secondary antibody (1:10,000 M.O.M. Biotinylated Anti Mouse Cat # MKB-2225-.1, Vector Laboratories Cat # BA-1000) was incubated for 2 hours with 20% Superblock (Fisher Scientific Cat # 37535). To enhance the signal intensity, tissues were incubated in an avidin-peroxidase solution (Vectastain Elite Cat # PK-6200). Subsequently, tissues were washed twice with 0.2 M sodium acetate, pH 6.0 for 10 min. Tissues were then transferred to the chromogen solution (0.3% hydrogen peroxide, 0.05% 3, 3-diaminobenzidine tetrahydrochloride, and 0.05% nickel ammonium sulfate hexahydrate in 0.2 M sodium acetate, pH 6). After achieving color development, tissues were washed twice with 0.2 M sodium acetate for 10 min and mounted from a PBS solution on positively charged microscope slides (Fisher Scientific Cat # 15-188-48). Once mounted and dried, slides underwent dehydration through Citrisolv and a series of alcohol gradients (70, 80, 95, and 100%) and were coverslipped (Fisher Scientific Cat 12545F) with DEPEX mounting medium (Electron Microscopy Sciences Cat # 13515).

### Imaging and quantification

The sections on slides were imaged at 20X using Olympus Slideview VS200 slide scanner microscope. The entire brain was used for digital quantification. Digital quantification was performed by adjusting the pixel classifier in QuPath (version −0.5.1) ^57^ to select positively stained pixels, and the percentage area above the threshold was quantified for analysis, blinded for the treatment group assignments. In addition, a blinded individual scored the slides in a semi quantitative manner using the entire brain.

### Statistics

The data obtained in **Figures 2-4, and 6** were subjected to statistical analysis using a one-way analysis of variance (ANOVA) test to evaluate the differences between groups. Following the ANOVA analysis, post hoc tests were conducted using Tukey’s Honestly Significant Difference (HSD) test to determine specific group differences. For the semi- and quantitative immunohistochemistry data (**Figure 7**), the non-parametric Kruskal-Wallis test was used to determine any significant group differences, with Dunn’s multiple comparisons test for the post hoc analyses to determine which groups were significantly different. The significance level was set at *p <* 0.05 to determine the statistical significance of the observed group differences. The statistical analyses of **Figures 2-4 and 6-7** were performed using GraphPad Prism 10. Gait statistical analyses (**Figure 5F-G**) were performed using IBM SPSS Statistics (version 28.0.0.1).

**Figure 3:**
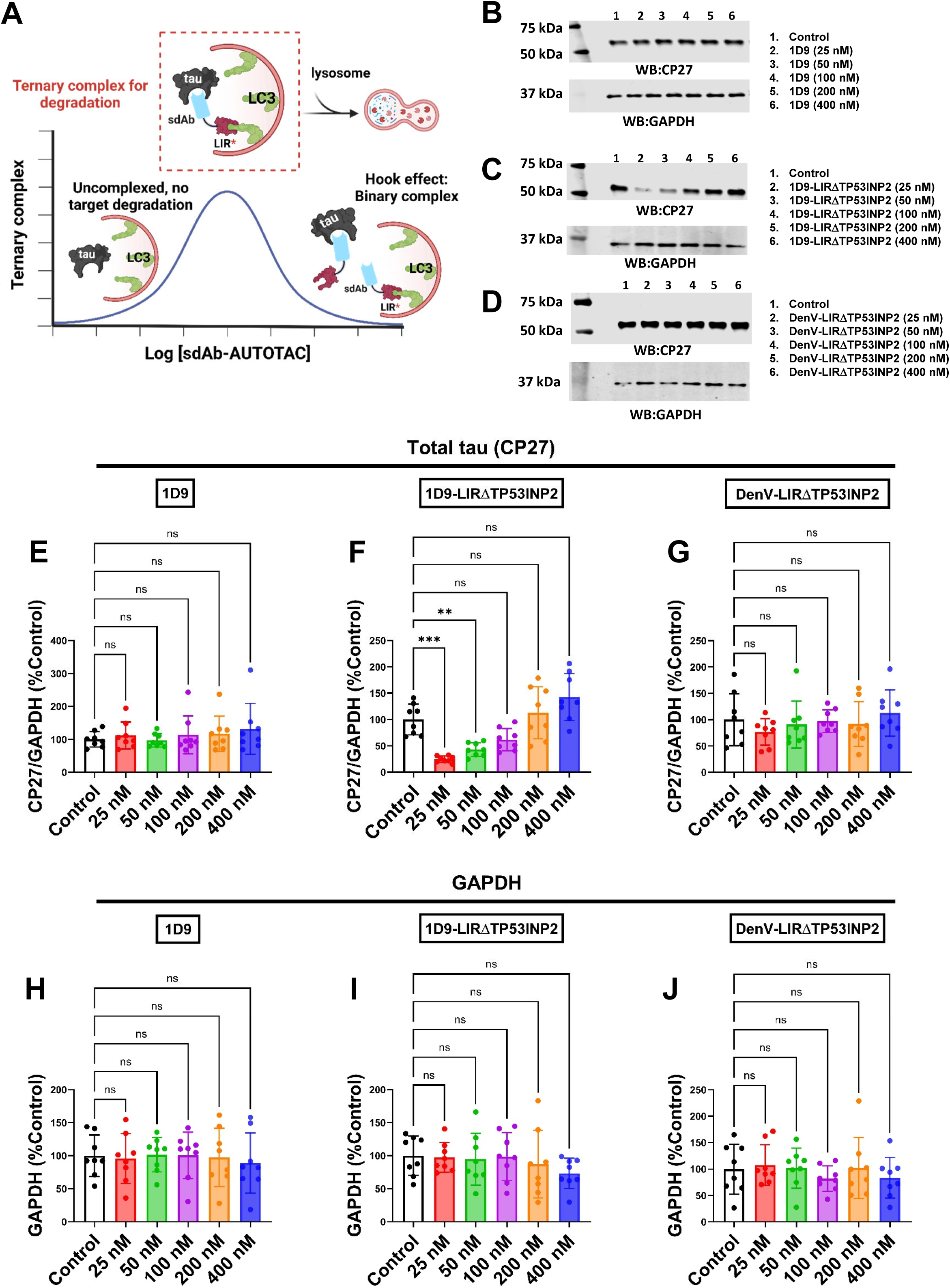
Kinetics of tau degradation in a human neuronal cell model of tauopathy. **(A)** Schematic diagram of single-domain antibody (sdAb)-based protein degrader-mediated ternary complex formation and hook effect at high concentrations. **(B-D)** Representative immunoblots depicting total tau (CP27) and GAPDH levels in treated human neuronal cell lysates treated with 1D9 (**B**), 1D9-LIRΔTP53INP2 (**C**), and control Denv-LIRΔTP53INP2 **(D)** Complete western blots and bands analyzed are in **Supplemental Figure 3**. Human neuronal cells were incubated with different concentrations of sdAbs ranging from 0–400 nM of either 1D9, 1D9-LIRΔTP53INP2, or control DenV-LIRΔTP53INP2 with samples collected 72 h post-antibody application (*n* = at least 8 replicates per condition). **(E-J)** One-way ANOVA analysis revealed no significant differences in normalized total tau levels (CP27) treated with various concentrations of 1D9 (*p* = 0.7636) (**E**). In contrast, 1D9-LIRΔTP53INP2 significantly decreased total tau levels (*p* < 0.0001) (**F**), whereas control DenV-LIRΔTP53INP2 did not (*p* = 0.6087) (**G**). (**F**) At the concentration of 25 nM and 50 nM, 1D9-LIRΔTP53INP2 led to 75% (*p* = 0.0003) and 57% (*p* = 0.0096) decrease in total tau levels (CP27) when compared to the control treatment. At the concentration of 100-400 nM, there were no significant differences between 1D9-LIRΔTP53INP2 and control treatments, indicating the presence of a hook effect. One-way ANOVA analysis revealed no significant differences in GAPDH levels treated with either 1D9 (*p* = 0.9869) (**H**), 1D9-LIRΔTP53INP2 (p = 0.6480) (**I**), or DenV-LIRΔTP53INP2 (*p* = 0.7555) (**J**). **, ***: *p* < 0.01, 0.001 (Tukey’s post-hoc test)

**Figure 4.**
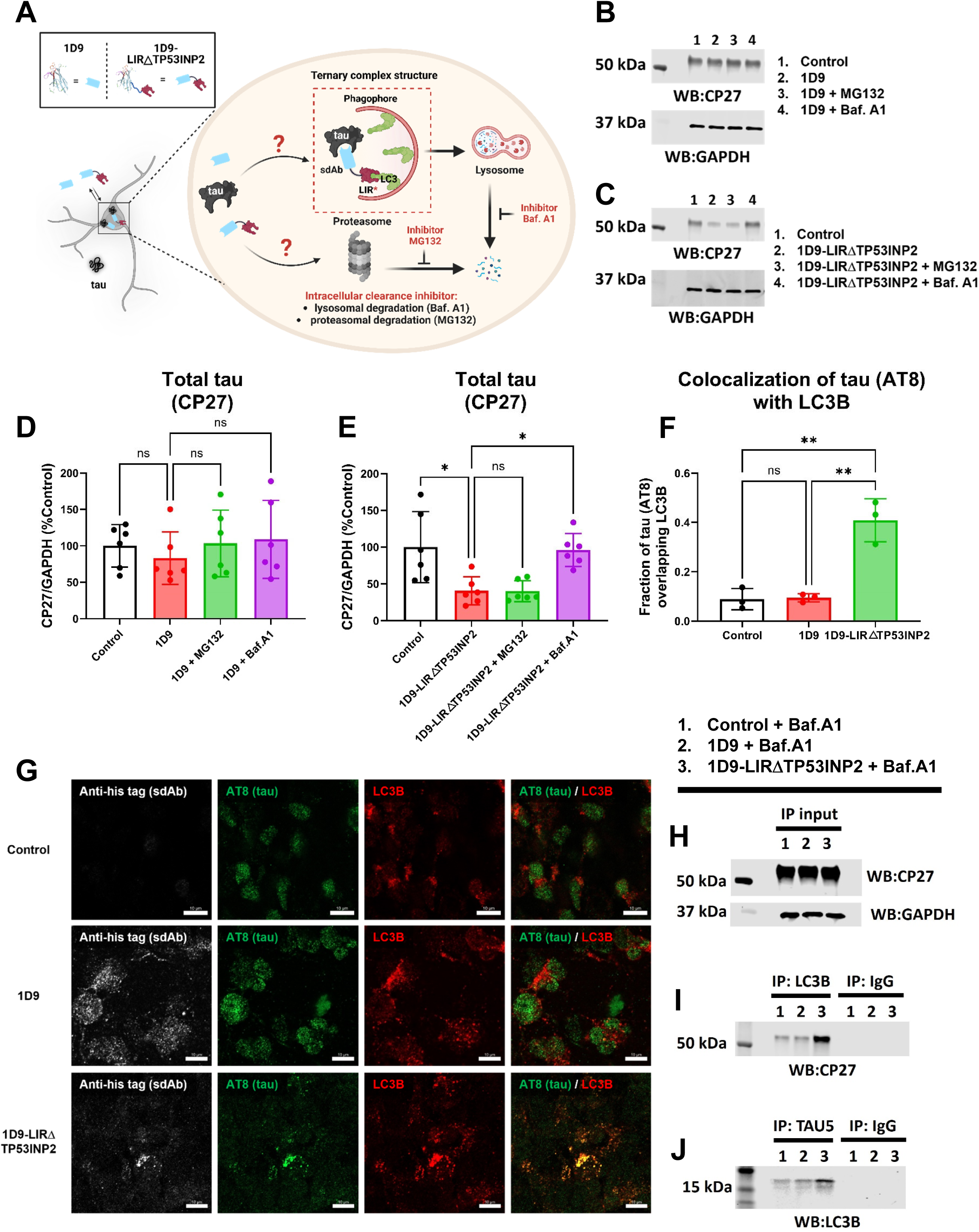
Mechanism of tau clearance by 1D9-LIRΔTP53INP2 via the autophagy-lysosome pathway in a human neuronal model of tauopathy. **(A)** To investigate the degradation mechanism, lysosomal and proteasomal pathways were blocked using bafilomycin A1 (Baf.A1, 0.2 µM) and MG132 (2 µM), respectively. P301L neurons were pretreated with inhibitors for 6 h, followed by treatment with vehicle, 25 nM 1D9, or 1D9-LIRΔTP53INP2 for 48 h (*n* = at least 6 replicates per condition). **(B-C)** Representative immunoblots of total tau (CP27) and GAPDH levels after treatment with 1D9 or 1D9-TP53INP2. **(D)** One-way ANOVA revealed no significant differences in normalized total tau levels (CP27) in the 1D9 treatment group (*p* = 0.7426). **(E)** One-way ANOVA showed significant differences in normalized total tau levels (CP27) in the 1D9-LIRΔTP53INP2 treatment group (*p* = 0.0012). 1D9-LIRΔTP53INP2 reduced total tau by 59% compared to vehicle control (*p* = 0.0104). This degradation was unaffected by MG132 (*p* > 0.9999) but was reversed by Baf.A1, increasing tau levels by 56% (*p* = 0.0172). **(F-G)** Quantification (**F**) and representative images (**G**) of sdAb (1D9 and 1D9-LIRΔTP53INP2), p-tau (AT8), and endogenous LC3B in P301L neurons. Cells were pretreated with Baf.A1 (0.2 µM) for 6 h, followed by treatment with vehicle, 1D9 or 1D9-LIRΔTP53INP2 for 24 h. Immunostaining for sdAb (His-tag, white), p-tau (AT8, green), and LC3B (red) were performed. Scale bar: 10 µm. Mander’s coefficient quantified tau-LC3B colocalization. One-way ANOVA indicated a significant increase in the fraction of p-tau overlapping with LC3B for 1D9-LIRΔTP53INP2 (*p* = 0.0007), with 357% (*p* = 0.0012) and 333% (*p* = 0.0013) increases compared to control and 1D9 groups, respectively. **(H-J)** Ternary complex formation in P301L neurons upon treatment with vehicle, 1D9, or 1D9-LIRΔTP53INP2, as analyzed by co-IP and western blot. Neurons were pretreated with Baf.A1 (0.2 µM) for 6 h, followed by 25 nM 1D9 or 1D9-LIRΔTP53INP2 for 48 h. **(H)** Western blot of 2% input. **(I)** LC3B pulldown and tau detection showed stronger tau bands in the 1D9-LIRΔTP53INP2 group. **(J)** Tau pulldown and LC3B detection revealed stronger LC3B bands in the 1D9-LIRΔTP53INP2 group. Complete western blot data are in **Supplemental Figure 4**. *, **: p < 0.05, 0.01, (Tukey’s post-hoc test).

**Figure 5:**
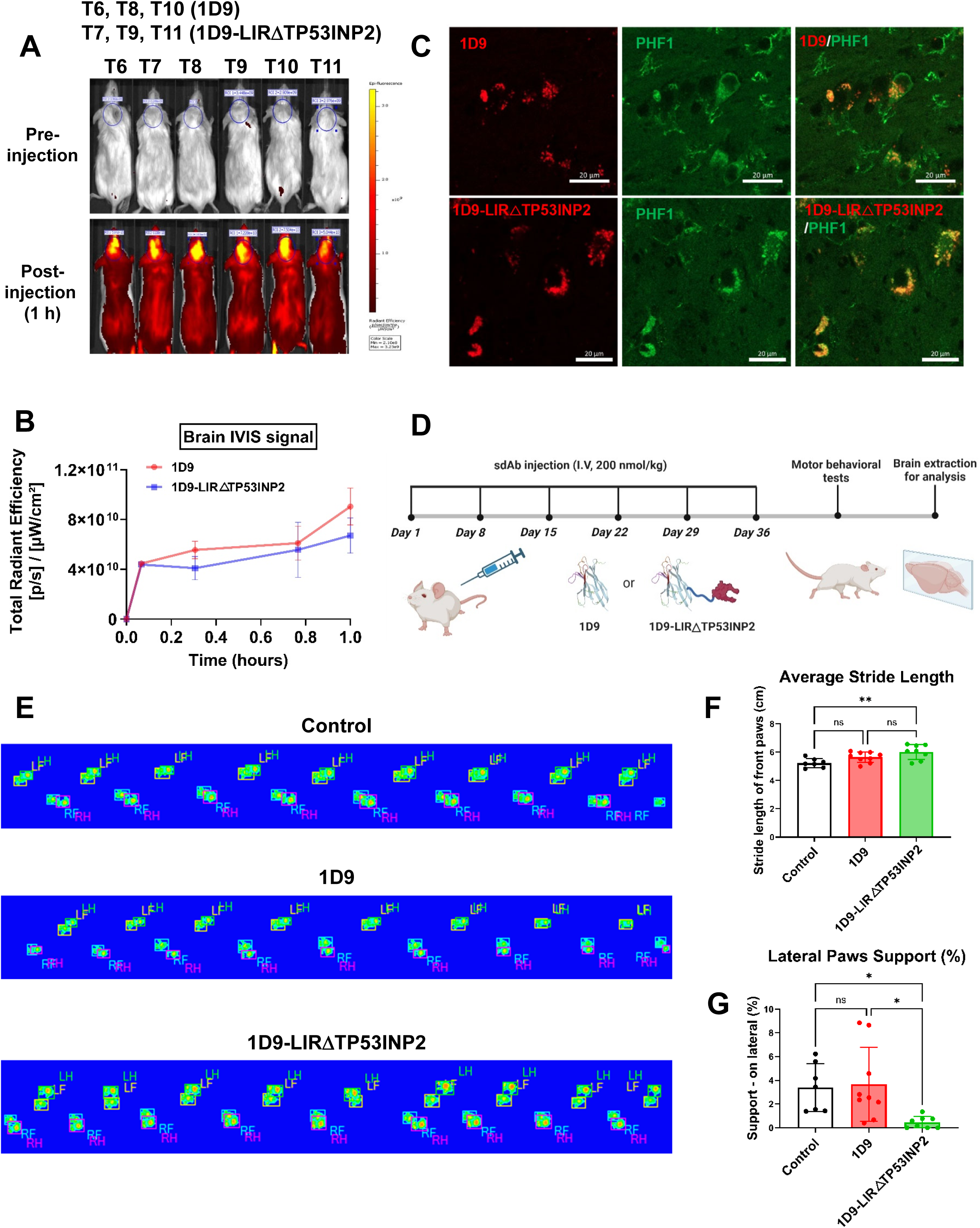
In-vivo efficacy of 1D9-LIRΔTP53INP2 in the JNPL3 mouse model of tauopathy. **(A)** Brain signal intensity after intravenous (i.v.) injection of near-infrared (NIR) tagged (680XL) sdAb-1D9 or 1D9-LIRΔTP53INP2 (700 nmol/kg), visualized by IVIS imaging. Scale bar indicates maximum pixel intensity. **(B)** Quantitative analysis of IVIS brain signal over time. **(C)** Colocalization of i.v. injected sdAb with p-tau (PHF1) in JNPL3 mice. NIR dye–labeled sdAb was injected i.v., and brains were perfused, sectioned, and stained with an antibody against p-tau (PHF1) 1-hour post-injection. Merged images confirm that 1D9 and 1D9-LIRΔTP53INP2 entered the brain, were taken up into neurons, and bound to pathological tau protein. Scale bars: 20 μm. Z-stack images are in **Supplemental Figure 5A**. **(D)** Schematic of the experimental protocol: JNPL3 mice received six weekly i.v. injections of vehicle, unmodified 1D9 (200 nmol/kg), or 1D9-LIRΔTP53INP2 (200 nmol/kg) in PBS. Following treatment, motor function was assessed with the CatWalk gait analysis system, and brains were collected for tissue analysis. **(E)** Representative CatWalk footprints from mice in each treatment group. **(F)** One-way ANOVA revealed significant differences in average stride length of four paws (*p* = 0.0052, *F* [2,21] = 6.841). Following 1D9-LIRΔTP53INP2 treatment, average stride length was increased by 15% (*p* = 0.0037) compared to the vehicle group. **(G)** One-way ANOVA revealed significant differences in the percentage of lateral paw support (**G**, *p* = 0.015, *F* [2,21] = 5.164). 1D9-LIRΔTP53INP2 reduced time spent in lateral support by 87% (*p* = 0.0473) compared to vehicle and by 88% (*p* = 0.0194) compared to 1D9 treatment. *, **, ***: p < 0.05, 0.01, 0.001, (Tukey’s post-hoc test).

**Figure 6:**
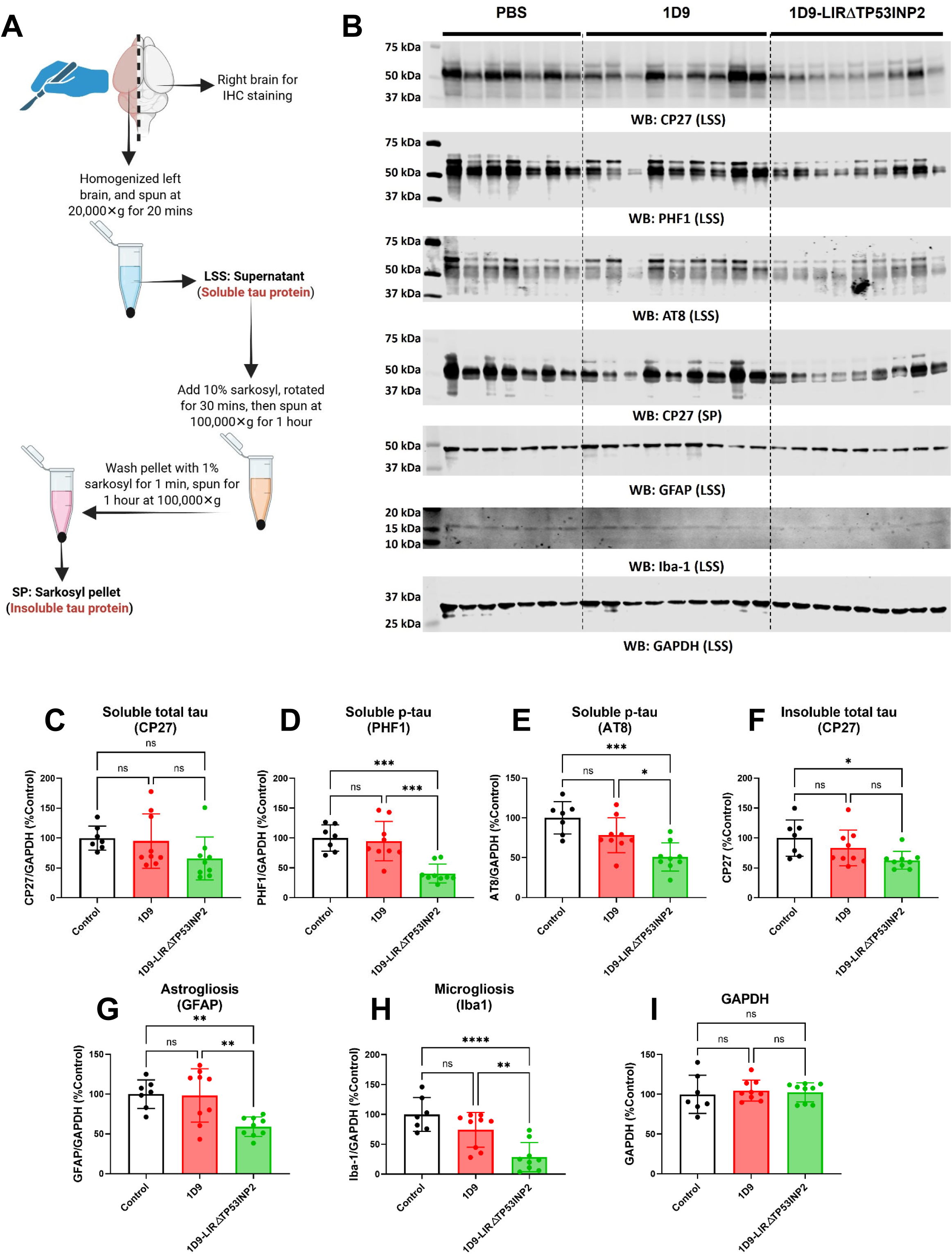
Western blot analysis of tau protein levels and gliosis markers in the in vivo study. **(A)** Schematic of the protocol for preparing soluble and insoluble tau fractions from mouse brain tissue. **(B)** Representative western blot showing soluble and insoluble tau levels alongside gliosis markers, probed with antibodies against total tau (CP27), p-tau (PHF1 and AT8), GFAP, Iba-1, and control GAPDH. **(C)** No significant differences in soluble total tau levels were observed between the groups (CP27, *p* = 0.1405; one-way ANOVA). **(D-E)** Significant group differences in soluble p-tau levels were observed with PHF1 (*p* < 0.0001, **D**) and AT8 staining (*p* = 0.0003, **E**). Soluble p-tau levels were significantly reduced following treatment with 1D9-LIRΔTP53INP2: PHF1 immunoreactive bands decreased by 60% compared to vehicle (*p* = 0.0003) and by 54% compared to unmodified 1D9 (*p* = 0.0004). AT8 immunoreactive bands decreased by 49% compared to vehicle (*p* = 0.0002) and by 27% compared to unmodified 1D9 (*p* = 0.0222). **(F)** Insoluble total tau levels (CP27) showed significant group differences (*p* = 0.027; one-way ANOVA), with 1D9-LIRΔTP53INP2 reducing total tau by 37% compared to vehicle (*p* = 0.0221). **(G-H)** Group differences were also noted in normalized GFAP (*p* = 0.002, **G**) and Iba-1 (*p* = 0.0001, **H**) levels. Treatment with 1D9-LIRΔTP53INP2 significantly reduced normalized gliosis marker levels: GFAP by 41% (*p* = 0.0061) and Iba-1 by 71% (*p* < 0.0001) compared to vehicle, and by 39% (*p* = 0.005) and 46% (*p* = 0.0048), respectively, compared to unmodified 1D9. \**p* < 0.05, \*\**p* < 0.01, \*\*\**p* < 0.001, \*\*\*\**p* < 0.0001 (Tukey’s post-hoc test). Full western blots and bands analyzed are provided in **Supplemental Figure 7.**

**Figure 7:**
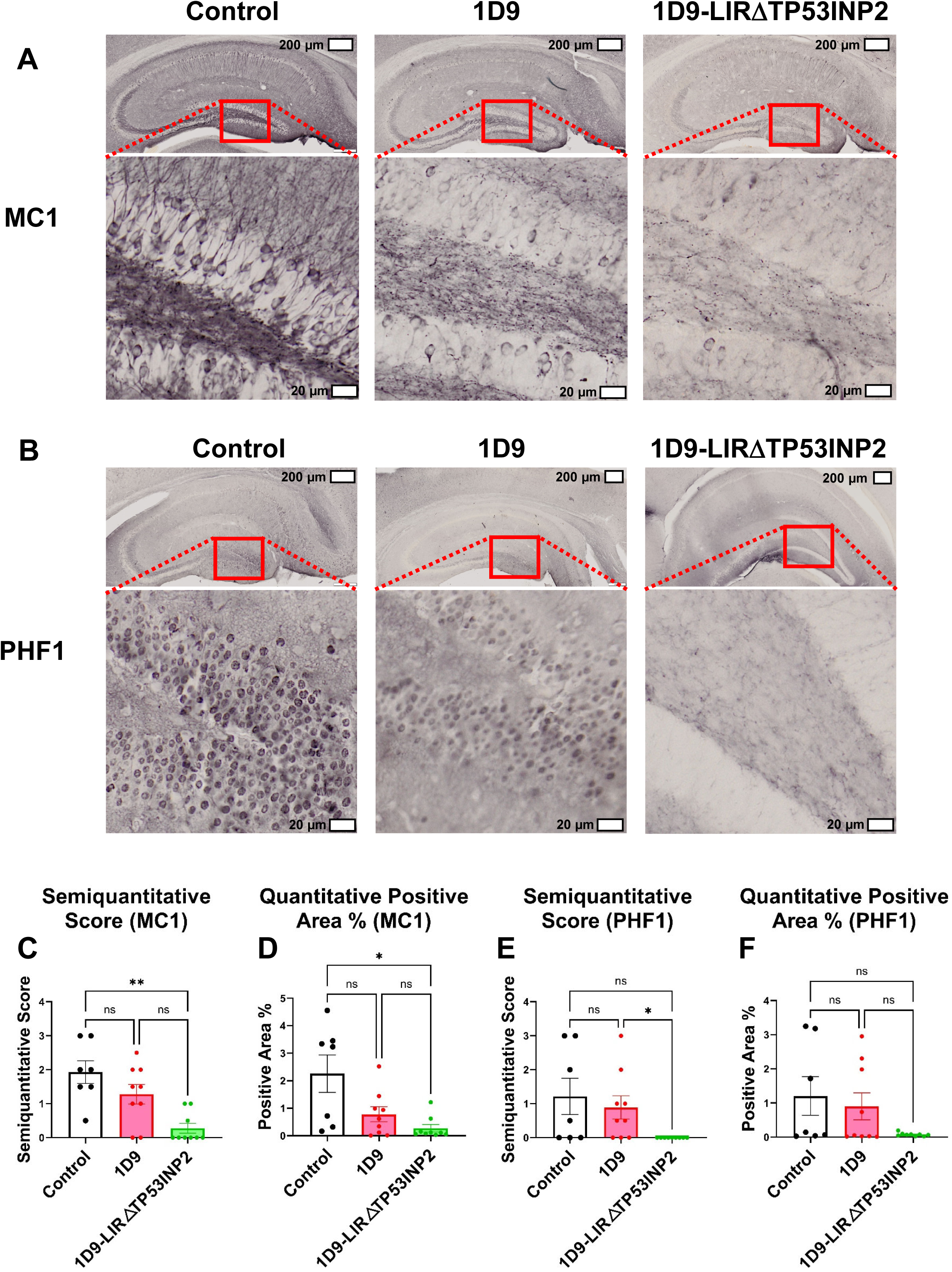
Effects of sdAb treatment on conformational (MC1) and phosphorylated (PHF1) tau pathology in JNPL3 mice. **(A-B)** Representative images of the hippocampus region from mice treated with PBS, 1D9, or 1D9-LIRΔTP53INP2, stained for (**A**) conformational tau (MC1) and (**B**) p-tau (PHF1). Upper panels show low magnification images (scale bars: 200 μm); red boxed regions are shown at higher magnification below (scale bars: 20 μm). (**C**) Semi-quantitative analysis of MC1-labeled tau pathology. Kruskal–Wallis test revealed significant group differences (*p* = 0.0028). Mice treated with 1D9-LIRΔTP53INP2 showed an 86% reduction in MC1 immunoreactivity compared to PBS-treated controls (*p* = 0.0025). No significant difference was observed between the 1D9 and 1D9-LIRΔTP53INP2 groups. (**D**) Quantitative analysis of MC1-labeled tau pathology. Kruskal–Wallis test showed significant differences among groups (*p* = 0.0203), with 88% reduction in tau pathology in the 1D9-LIRΔTP53INP2 group relative to PBS (*p* = 0.016). No significant difference was found between 1D9 and 1D9-LIRΔTP53INP2. (**E**) Semi-quantitative analysis of PHF1-labeled tau pathology. Group differences were significant (Kruskal Wallis, *p* = 0.0157), with a complete (100%) reduction in PHF1 signal in the 1D9-LIRΔTP53INP2 group compared to the 1D9 group (*p* = 0.0318). (**F**) Quantitative analysis of PHF1-labeled tau pathology. No significant differences were observed among treatment groups. \**p* < 0.05, \*\**p* < 0.01, (Kruskal-Wallis test).

## Results

### Design and in vitro evaluation of targeted tau degraders

We have recently developed two promising anti-tau single-domain antibody (sdAb) candidates, 2B8 and 1D9, which exhibit therapeutic and diagnostic potential in primary culture model and mouse model of tauopathies ^15, 17^. Our phage display library, constructed from B-cells of a llama immunized with recombinant human tau (2N4R) and enriched paired helical filament (PHF) tau isolated from human tauopathy brain tissue, yielded 58 unique anti-tau sdAb clones. Following iterative screening for binding affinity to various tau preparations and evaluating efficacy in a tauopathy primary culture model, while also factoring in solubility for antibody engineering, we selected anti-tau sdAb 1D9 for further antibody engineering development.

To amplify therapeutic efficacy, we engineered anti-tau sdAb 1D9 by fusing an LC3-interacting region (LIR) motif via peptide linkers. This strategy integrates sdAb-based immunotherapy with intracellular targeted degradation approaches within the autophagy-lysosome pathway ^58^. Our objective is to determine whether the engineered derivative of 1D9 exhibits improved efficacy compared to its non-engineered counterpart.

To this end, we developed two engineered derivatives of 1D9 with differing lengths of LIR motifs and conjugation linkers. The shorter version 1D9-LIRΔp62 linked the sdAb 1D9 to a shorter LIR peptide motif derived from the autophagy receptor p62 (residues 332–343: SGGDDDWTHLSS ^59^) using a generic linker (GSGS) (**Figure 1C**). In the prior study ^59, 60^, this shorter LIR peptide motif was conjugated to an α-synuclein-targeting peptide to create an α-synuclein degrader that utilizes the autophagy-lysosome pathway. However, the α-synuclein-binding peptide likely has weaker affinity and lower in vivo stability compared to the sdAb.

In contrast, the longer version 1D9-LIRΔTP53INP2 was conjugated to the sdAb 1D9 with a mutant LIR motif derived from the Tumor Protein 53 Induced Nuclear Protein 2 (TP53INP2), known for its enhanced recognition of the LC3 protein ^61^. This mutant motif was previously linked to an anti-Alfa tag sdAb through an extended peptide linker (SGGSSGGSSGSETPGTSESATPESSGGSSGGS), facilitating the efficient degradation of Alfa tag-labeled aggregation-prone proteins including tau and α-synuclein in co-transfected human osteosarcoma (U2OS) cells, as well as tagged organelles ^62^. However, Alfa tag-labeled proteins and organelles may not fully recapitulate the native disease state, and U2OS cells—being derived from cancer—may not adequately model neurodegenerative conditions.

Additionally, we generated a corresponding negative control sdAb chimera DenV-LIRΔTP53INP2 by substituting the anti-tau sdAb 1D9 with an anti-Dengue virus (DenV) sdAb, attached to the extended linker and LIR motif, effectively abrogating tau-binding capacity (**Figure 1C**).

We first assessed the impact of antibody engineering on the antigen recognition ability of sdAb 1D9. A dot-blot assay was performed using two tau antigens—recombinant hyperphosphorylated tau (rec p-tau) and patient-derived paired helical filament (PHF) enriched tau — along with a negative control, bovine serum albumin (BSA). The analysis demonstrated no significant differences in antigen binding among 1D9, 1D9-LIRΔp62, and 1D9-LIRΔTP53INP2 in both the rec p-tau and PHF-tau groups, indicating that these antibody engineering modifications do not impair the ability of sdAb 1D9 to recognize tau (**Figure 1D and Supplemental Figure 1A**).

In contrast, DenV-LIRΔTP53INP2 showed no detectable signal compared to 1D9 (rec p-tau: *p* = 0.0007; PHF-tau: *p* = 0.0073), 1D9-LIRΔp62 (rec p-tau: *p* = 0.0497; PHF-tau: *p* = 0.0302), and 1D9-LIRΔTP53INP2 (rec p-tau: *p* = 0.0116; PHF-tau: *p* = 0.0005) in both rec p-tau and PHF-tau groups. These findings indicate that substituting the anti-tau sdAb 1D9 with an anti-Dengue virus (DenV) sdAb abolishes tau-binding capability (**Figure 1D and Supplemental Figure 1A**). No significant signals were observed in the BSA group, serving as a negative control.

Furthermore, we employed biolayer interferometry (BLI) to quantify the in vitro binding affinity of the sdAb degrader molecule towards the same two tau variants as for the dot blot assay: rec p-tau and PHF-tau. The non-engineered sdAb 1D9 exhibited binding affinities of 30.3 nM and 84.4 nM towards rec p-tau and the PHF-tau, respectively. The engineered sdAbs (1D9-LIRΔp62 and 1D9-LIRΔTP53INP2) demonstrated comparable binding to these tau variants, with affinities ranging from 32.0 to 36.7 nM for rec p-tau and 63.6 to 76.1 nM for the PHF-tau (**Figure 1E and Supplemental Figure 1B-G**). Both non-engineered and engineered sdAbs showed similar association (k_a_) and dissociation (k_d_) rates when interacting with rec p-tau and PHF-tau, indicating that the modifications did not significantly alter sdAbs’ affinities towards the tau antigens (**Figure 1E**).

### Evaluating the degradation efficacy of sdAbs targeting tau in a human neuronal cell model of tauopathy

To assess tau degradation efficacy of sdAbs, we utilized a patient-derived induced pluripotent stem cell (iPSC) model, which is increasingly recognized as a robust tool for exploring the molecular mechanisms underlying neurodegenerative diseases and for developing potential therapeutic strategies ^39, 54^. In this study, we derived neurons from an individual with frontotemporal dementia (FTD) who carries the tau-P301L autosomal dominant mutation. Post-mitotic neurons were generated from cortical-enriched neural progenitor cells (NPCs) derived from the patient’s iPSCs, which were differentiated into mature cortical neurons over 4 to 6 weeks, as previously reported ^54^ (**Figure 2A**). These patient-specific neuronal models express tau at endogenous levels, accurately reflecting disease-relevant tau characteristics in vitro. Therefore, these models provide a valuable platform for evaluating therapeutic candidates in a clinically relevant human disease context.

After differentiating NPCs into P301L neurons, we treated them with sdAbs at a concentration of 50 nM for 3 days (**Figure 2A**), compared to untreated control, followed by western blot analysis to evaluate intracellular tau clearance. One-way ANOVA revealed significant differences in normalized total tau levels (CP27, *p* < 0.0001) and p-tau levels (PHF-1, *p* < 0.0001; pSer199, *p* < 0.0001). Treatment with 1D9-LIRΔTP53INP2 resulted in a 41% reduction in total tau levels (CP27, *p* = 0.0011). Notably, 1D9-LIRΔTP53INP2 exhibited a more pronounced degradation effect on p-tau, with reductions of 82% for PHF1 and 68% for pSer199 (*p* < 0.0001 for both), compared to the vehicle control group (**Figure 2B-D, G**). These results suggest that 1D9-LIRΔTP53INP2 may selectively degrade pathological p-tau relative to physiological total tau levels. This selectivity may stem from the design of 1D9-LIRΔTP53INP2, which aims to enhance tau degradation by bringing tau and autophagic machinery into proximity. It is hypothesized that larger, aggregated proteins are more likely to be degraded via the autophagy-lysosome pathway, while smaller soluble proteins are typically processed through proteasomal degradation ^44^.

Moreover, 1D9-LIRΔTP53INP2 demonstrated significantly greater degradation of both total and p-tau compared to treatments with 1D9 and 1D9-LIRΔp62. Specifically, 1D9-LIRΔTP53INP2 led to a 43% reduction in total tau (CP27, *p* = 0.0005) and 67% and 49% reductions in PHF1 and pSer199 immunoreactive tau, respectively, compared to the 1D9 treatment group (PHF1, *p* < 0.0001; pSer199, *p* = 0.0006). When compared to the 1D9-LIRΔp62 treatment group, 1D9-LIRΔTP53INP2 achieved a 49% reduction in total tau (CP27, *p* < 0.0001) and reductions of 81% and 46% in PHF1 and pSer199 immunoreactive tau, respectively (PHF1, *p* < 0.0001; pSer199, *p* = 0.0014). Importantly, one-way ANOVA indicated no significant differences in GAPDH (*p* = 0.2706) or NeuN levels (*p* = 0.9283), indicating that neither unmodified sdAb 1D9 nor the two modified sdAb 1D9 chimeras exhibit toxicity at a concentration of 50 nM (**Figure 2E-G**).

### Kinetics of tau degradation

The hook effect is frequently observed in hetero-bifunctional compounds designed for specific protein degradation, such as PROteolysis TArgeting Chimeras (PROTACs)^63^. This phenomenon occurs because the degradation mechanism relies on the formation of a ternary complex among the target protein, a specific ligase, and the hetero-bifunctional compound. At high compound concentrations, unproductive binary complexes form, rather than the productive trimeric complex necessary for degradation (**Figure 3A**). Similar phenomena have also been observed in the autophagosome-anchoring targeting chimera (AUTOTAC) technique using small molecules to degrade tau protein ^50^, including the 1D9-AUTOTAC derivatives examined here.

To establish the kinetics of tau degradation, we treated P301L neurons with five different sdAb concentrations ranging from 0 to 400 nM for 72 hours. One-way ANOVA revealed no significant differences in normalized total tau levels (CP27, *p* = 0.7636) or GAPDH levels (*p* = 0.9869) for the 1D9 treatment, indicating that 1D9 showed neither tau degradation effect nor toxicity under these conditions (**Figure 3B, E, H**). However, we have observed a tau degradation effect of 1D9 sdAb at a concentration of 1 µg/ml (∼50 nM) in JNPL3 tauopathy primary cultures treated with patient-derived paired helical filament (PHF)-enriched tau ^17^.

In the 1D9-LIRΔTP53INP2 treatment group, one-way ANOVA revealed significant differences in normalized total tau levels (CP27, *p* < 0.0001). At concentrations of 25 nM and 50 nM, 1D9-LIRΔTP53INP2 showed a 75% (*p* = 0.0003) and 57% (*p* = 0.0096) reduction in total tau levels (CP27) compared to the vehicle control treatment. However, at concentrations higher than 100 nM, no significant differences were observed between the 1D9-LIRΔTP53INP2 and control treatments, indicating the presence of hook effects at higher concentrations (**Figure 3A, C, F**). One-way ANOVA also revealed no significant differences in GAPDH levels (*p* = 0.6480), indicating lack of toxicity (**Figure 3C, I**).

Conversely, we tested the tau degradation efficacy of the negative control DenV-LIRΔTP53INP2. One-way ANOVA revealed no significant differences in normalized total tau levels (CP27, *p* = 0.6087) or GAPDH levels (*p* = 0.7555) (**Figure 3D, G, J**). Given the lack of tau-binding capacity due to replacing the anti-tau 1D9 sdAb, DenV-LIRΔTP53INP2 did not show any tau degradation efficacy at concentrations ranging from 0 to 400 nM. This indicates that engagement with tau via anti-tau sdAb 1D9 is required for 1D9-LIRΔTP53INP2-mediated tau clearance (**Figure 1B**).

### Degradation pathways of sdAbs targeting tau in a human neuronal cell model of tauopathy

To investigate the mechanism of action of 1D9-LIRΔTP53INP2, we focused on two prominent degradation pathways: the lysosomal and proteasomal pathways. We employed the lysosomal inhibitor bafilomycin A1 (Baf.A1) and the proteasomal inhibitor MG132 to selectively inhibit these pathways (**Figure 4A**). P301L neurons were pre-treated for 6 hours with either Baf.A1 (0.2 µM) or MG132 (2 µM) before a subsequent 2-day treatment with 25 nM of either 1D9 or 1D9-LIRΔTP53INP2, followed by western blot analysis.

In the 1D9 treatment group, one-way ANOVA indicated no significant difference in normalized total tau levels (CP27, *p* = 0.7426) (**Figure 4B, D**). Conversely, in the 1D9-LIRΔTP53INP2 treatment group, a significant difference was observed in normalized total tau levels (CP27, *p* = 0.0012). Notably, 1D9-LIRΔTP53INP2 without inhibitors resulted in a 59% reduction in total tau levels (CP27, *p* = 0.0104) compared to the vehicle control. The application of the proteasomal inhibitor MG132 did not reverse this degradation effect (*p* > 0.9999); however, the lysosomal inhibitor Baf.A1 significantly mitigated the degradation, increasing normalized total tau levels (CP27) by 56% (*p* = 0.0172) (**Figure 4C, E**). These findings indicate that 1D9-LIRΔTP53INP2 facilitates tau protein degradation primarily through the lysosomal pathway.

Additionally, we assessed whether the degradation mediated by 1D9-LIRΔTP53INP2 involved the autophagy-lysosome process. To this end, we performed immunofluorescence (IF) to examine the cellular localization of the sdAb, tau, and endogenous autophagy marker LC3B protein. P301L neurons were pre-treated with Baf.A1 (0.2 µM) for 6 hours, followed by a 24-hour treatment with 25 nM of either 1D9 or 1D9-LIRΔTP53INP2, and subsequently analyzed via immunofluorescence.

In the control and sdAb 1D9 treatment, p-tau protein (AT8) appeared as amorphous aggregates dispersed throughout the cytoplasm, with minimal colocalization with LC3B (**Figure 4F-G**). In contrast, treatment with 1D9-LIRΔTP53INP2 resulted in a morphological transformation of p-tau (AT8) into punctate structures within the cytoplasm, demonstrating significant colocalization with both 1D9-LIRΔTP53INP2 and endogenous LC3B (**Figure 4G**). One-way ANOVA revealed a significant difference in the fraction of tau (AT8) overlapping with endogenous LC3B (*p* = 0.0007). Specifically, 1D9-LIRΔTP53INP2 treatment resulted in a 357% (*p* = 0.0012) increase in colocalization compared to the control group and a 333% (*p* = 0.0013) increase compared to the 1D9 treatment group (**Figure 4F**).

To address potential false positives in colocalization due to overlapping proteins in different layers during confocal imaging, we conducted co-immunoprecipitation (co-IP) assays to further investigate the interactions among 1D9-LIRΔTP53INP2, tau, and LC3B at the molecular level. P301L neurons were pre-treated for 6 hours with the lysosomal inhibitor Baf.A1 (0.2 µM) to promote the accumulation of ternary complexes prior to tau clearance, followed by a 48-hour treatment with 25 nM of either 1D9 or 1D9-LIRΔTP53INP2, and subsequent co-IP assays to explore ternary complex interactions (**Figure 4A**).

Immunoprecipitation of LC3B and subsequent western blot analysis revealed co-IP with total tau (CP27). A weak signal for total tau was detected in the vehicle control and 1D9 treatment groups, suggesting some endogenous interaction between tau and LC3B in human neurons. Conversely, a stronger total tau band was observed in the 1D9-LIRΔTP53INP2 treatment group, indicating enhanced interaction between tau and LC3B initiated by 1D9-LIRΔTP53INP2 treatment (**Figure 4I**). Similarly, when immunoprecipitating total tau, western blot analysis showed co-IP with LC3B. Weak detection of LC3B was noted in the vehicle control and 1D9 treatment groups, while a stronger band was evident in the 1D9-LIRΔTP53INP2 treatment group (**Figure 4J**). These results support the formation of a ternary complex among 1D9-LIRΔTP53INP2, tau, and LC3B, providing further evidence for the mechanism of tau clearance by 1D9-LIRΔTP53INP2 through the function of LC3B and lysosomal degradation.

### Degradation efficacy of sdAbs targeting tau in the JNPL3 mouse model of tauopathy

In the next phase of our study, we evaluated the in vivo effectiveness of the unmodified sdAb 1D9 and the modified sdAb 1D9-LIRΔTP53INP2 in a JNPL3 mouse model of tauopathy with the P301L tau mutation. Prior to the in vivo experiment, we aimed to determine whether both sdAbs could traverse the blood-brain barrier (BBB) and reach intraneuronal tau aggregates. To this end, we labeled the unmodified sdAb 1D9 and the modified sdAb 1D9-LIRΔTP53INP2 with a near-infrared (NIR) tag, VivoTag 680XL ^15^.

Homozygous tauopathy JNPL3 mice, 8 months of age, received a single intravenous injection of either NIR-labeled 1D9 or 1D9-LIRΔTP53INP2 (700 nmol/kg, ∼11.1 mg/kg for 1D9 and ∼16.9 mg/kg for 1D9-LIRΔTP53INP2). Post-injection, the NIR signal in the brain region increased gradually over 60 minutes (indicated by yellow in the region of interest, ROI, circle). The signal was detected throughout the body, with maximum intensity observed in the brain. The brain signal coincides with signal from the spinal cord, which also exhibits extensive tau pathology in the JNPL3 model (**Figure 5A-B**). One hour following the intravenous injection, brains were extracted, and the right hemisphere was immersion-fixed in paraformaldehyde, processed, sectioned, and stained with tau antibodies against p-tau (PHF1) and markers of autophagosomes (LC3B).

Confocal results indicated that both sdAb 1D9 and 1D9-LIRΔTP53INP2 were effectively detected in the brain, where they colocalized with pathological p-tau (PHF1) and autophagosome markers (LC3B) (**Figure 5C, Supplemental Figure 5A-C**). Notably, similar to the observations from human neuronal cultures (**Figure 4G**), the 1D9 treatment group exhibited p-tau as diffuse, amorphous aggregates throughout the cytoplasm. In contrast, the 1D9-LIRΔTP53INP2 group showed a distinct morphology, with p-tau appearing as punctate aggregates within the cytoplasm (**Figure 5C, Supplemental Figure 5A**). These findings suggest that following injection, 1D9-LIRΔTP53INP2 may bind to p-tau, directing it to autophagosomes (LC3B) for degradation. This hypothesis is also corroborated by the prominent bands observed in Co-IP assays, indicating the formation of a ternary complex among 1D9-LIRΔTP53INP2, tau, and LC3B (**Figure 4I-J**).

Following confirmation of BBB penetration and the engagement of intraneuronal tau aggregates by the sdAb, we conducted an in vivo study. We enrolled 25 female homozygous JNPL3 mice, aged 6-7 months, and divided them into three groups: vehicle control (*n*=7), 1D9 (*n*=9), and 1D9-LIRΔTP53INP2 (*n*=9). Each group received six intravenous injections—administered weekly—of either the vehicle control, unmodified sdAb 1D9 (200 nmol/kg, ∼3.2 mg/kg), or modified sdAb 1D9-LIRΔTP53INP2 (200 nmol/kg, ∼4.9 mg/kg) in PBS. Following antibody administration, we conducted Catwalk^TM^ locomotor analysis, which enables non-invasive evaluation of motor behavior ^64^. JNPL3 have motor impairments as originally described ^55^, and confirmed by us in several studies ^65, 66^, although the severity of this deficit has over the years shifted to older age. Subsequently, the mice were perfused, and their brains were extracted for tissue analysis (**Figure 5D**).

To better contextualize the effects of antibody treatment, we first conducted CatWalk™ gait analysis on a baseline cohort of untreated wild-type (*n* = 7) and JNPL3 (*n* = 9) female mice at 8-9 months of age—the same age as the treated cohort. Consistent with previous findings in P301S tauopathy mice ^67^, we observed that average stride length – the distance between successive placements of the same paw – was 9.8% shorter in JNPL3 tauopathy mice compared to wild-type mice across all four paws (*F* [1,15] = 5.715, *p* = 0.0314; **Supplemental Figure 6A**). This result was significant even when average speed was included as a covariate in the analysis (*F* [1,15] = 5.600, *p* = 0.034). We also observed that JNPL3 mice exhibited a 124% increase in the percentage of lateral paw support relative to wild-type mice (*F* [3,23] = 3.301, *p* = 0.041; **Supplemental Figure 6B**), where average speed had no significant impact when included as covariate. Increased reliance on lateral support is indicative of impaired gait stability, consistent with findings in stroke ^68^ and amyotrophic lateral sclerosis (ALS) ^69^ models.

We then assessed a second cohort of 8–9-month-old JNPL3 female mice following antibody treatment (PBS: *n*=7; 1D9: *n*=9; 1D9-LIRΔTP53INP2: *n*=9) using the Noldus CatWalk™ system. Representative footprint images for each treatment group are shown in **Figure 5E**. One-way ANOVA revealed a significant effect of treatment on average stride length of all four paws (*F* [2,21] = 6.841, *p* = 0.0052). Specifically, treatment with 1D9-LIRΔTP53INP2 increased average stride length by 15% compared to the vehicle control group (*p* = 0.0037; **Figure 5F**). Average speed (*F* [1,23] = 2.566, *p* = 0.126) and body weight (*F* [1,23] = 2.676, *p* = 0.118) were not significant as covariates in the analysis of stride length.

In the treated JNPL3 cohort, one-way ANOVA indicated significant differences across groups in the percentage of lateral support (*F* [2,21] = 5.164, *p* = 0.015). Notably, 1D9-LIRΔTP53INP2 treatment reduced lateral support time by 87% compared to vehicle (*p* = 0.0473) and by 88% compared to the unmodified 1D9 group (*p* = 0.0194; **Figure 5G**). Again, average speed (*F* [1,23] = 0.160, *p* = 0.694) and body weight (*F* [1,23] = 1.159, *p* = 0.295) were not significant as covariates in the analysis of lateral support. These findings demonstrate that 1D9-LIRΔTP53INP2 treatment enhances gait stability and motor function in JNPL3 mice.

We next investigated whether the observed motor function improvements correlated with reductions in tau levels and gliosis in the brain. After behavioral assessments, mice were perfused, and their brains were collected for analysis. The left hemisphere was homogenized to isolate soluble and insoluble tau protein fractions, while the right hemisphere was coronally sectioned into 40 μm sections for immunohistochemistry (**Figure 6A**).

For soluble total tau protein detected using the CP27 antibody, 1D9-LIRΔTP53INP2 treatment showed a trend toward reduced tau levels compared to both the control and unmodified 1D9 treatment groups, though this difference did not reach statistical significance (**Figure 6B-C**, one-way ANOVA, *p* = 0.1405). This observation aligns with previous findings in human neuronal cultures, where 1D9-LIRΔTP53INP2 selectively degraded pathological p-tau over physiological total tau (**Figure 2B-D**). In contrast, 1D9-LIRΔTP53INP2 significantly reduced pathological p-tau. Immunoblots with PHF1 (pSer396/404) and AT8 (pSer202/Thr205) antibodies revealed significant group differences for soluble p-tau levels (PHF1, *p* < 0.0001; AT8, *p* = 0.0003; one-way ANOVA) (**Figure 6B, D-E**). Specifically, 1D9-LIRΔTP53INP2 treatment reduced soluble PHF1 and AT8 immunoreactive p-tau levels by 60% and 49%, respectively, compared to the vehicle control group (PHF1, *p* = 0.0003; AT8, *p* = 0.0002), and by 54% and 27%, respectively, compared to the unmodified 1D9 group (PHF1, *p* = 0.0004; AT8, *p* = 0.0222). For total insoluble tau, significant group differences were also observed (**Figure 6B, F**, *p* = 0.027, one-way ANOVA), with 1D9-LIRΔTP53INP2 reducing levels by 37% compared to the vehicle control (CP27, *p* = 0.0221), though this reduction was not significant compared to the unmodified 1D9 group (*p* = 0.2169). In summary, in the JNPL3 tauopathy mouse model, 1D9-LIRΔTP53INP2 significantly reduced soluble pathological p-tau compared to both the control and 1D9 treatment groups, without affecting soluble total tau levels. Furthermore, it significantly reduced insoluble total tau levels compared to the control group.

Given the close association between tauopathy pathogenesis and glial activation ^70, 71^ and the presence of astrocytosis and microgliosis in the JNPL3 mice ^72^, we also evaluated whether 1D9-LIRΔTP53INP2 treatment could attenuate immune responses in glial cells. To assess this, we measured GFAP and Iba1 levels as markers of astrocytic and microglial activation, respectively, in treated mouse brains. Western blot analyses were conducted using the low-speed supernatant (LSS) fraction of brain homogenates. Normalized levels of GFAP (**Figure 6B, G**) and Iba-1 (**Figure 6B, H**) showed significant differences among the treatment groups (GFAP, *p* = 0.002; Iba-1, *p* = 0.0001; one-way ANOVA). Specifically, 1D9-LIRΔTP53INP2 treatment led to a 41% reduction in GFAP (*p* = 0.0061) and a 71% reduction in Iba-1 (*p* < 0.0001) compared to the vehicle control group. Compared to the unmodified 1D9 treatment, 1D9-LIRΔTP53INP2 reduced GFAP and Iba-1 levels by 39% (*p* = 0.005) and 46% (*p* = 0.0048), respectively. These results indicate that 1D9-LIRΔTP53INP2-mediated tau clearance effectively reduces gliosis in the brain.

GAPDH levels did not differ between the groups (**Figure 6B** and I; one-way ANOVA, p = 0.8542), indicating the absence of treatment-related toxicity.

Subsequently, we performed immunohistochemical analysis on the right hemisphere of mouse brains using conformational tau (MC1) and p-tau (PHF1) antibodies ^26^, as described in the Methods section. In the PBS-treated control group, conformational (MC1) and p-tau (PHF1) aggregates were detected throughout the brain. Notably, treatment with 1D9-LIRΔTP53INP2 markedly reduced MC1-positive conformational and p-tau (PHF1) aggregates compared to the control group (**Figure 7A-B**)

The immunohistochemical results were evaluated by both semi-quantitative and quantitative analyses (**Figure 7C-F**). For the MC1, semi-quantitative analysis revealed significant group differences (Kruskal-Wallis test, *p* = 0.0028), with 1D9-LIRΔTP53INP2 reducing conformational tau staining by 86% (*p* = 0.0025) (**Figure 7C**). Similarly, quantitative analysis confirmed significant differences (Kruskal-Wallis test, *p* = 0.0203), showing an 88% reduction in MC1 immunoreactivity (*p* = 0.016) (**Figure 7D**). There was no significant difference between the 1D9 and 1D9-LIRΔTP53INP2 treatment groups in either analysis.

For p-tau (PHF1) aggregates, significant group differences were observed only in the semi-quantitative analysis (Kruskal-Wallis test, *p* = 0.0157), with 1D9-LIRΔTP53INP2 reducing PHF1 immunoreactivity by 100% compared to the 1D9 group (*p* = 0.0318) (**Figure 7E**), although a similar trend was seen in the quantitative analysis but did not reach statistical significance (**Figure 7F**).

Together, these findings provide insight into the mechanisms underlying sdAb-mediated tau clearance and support the therapeutic potential of the engineered sdAb 1D9-LIRΔTP53INP2.

## Discussion

Intracellular protein aggregates are a hallmark of many neurodegenerative diseases, yet no approved therapies directly target them. Given that the majority of pathological aggregates form within cells, effective therapeutic strategies should be able to work intracellularly as we have discussed previously ^12^.

Currently, most tau-targeting agents in clinical trials are immunotherapy. However, these therapies primarily rely on full-length IgG antibodies, which have limited brain penetration, potentially reducing their effectiveness ^1^. In contrast, sdAbs, due to their smaller size (15 kDa vs. 150 kDa), exhibit improved brain entry, better tissue diffusion, and the ability to bind cryptic epitopes inaccessible to conventional antibodies. This advantage extends beyond tauopathy, offering potential for immunotherapies targeting other neurodegenerative diseases.

Additionally, sdAbs exhibit lower immunogenicity than whole antibodies, potentially leading to improved safety outcomes. Lessons from anti-amyloid-β (Aβ) antibody therapies, such as lecanemab ^73^ and donanemab ^74^, provide valuable insights. These antibodies have shown modest but significant cognitive benefits in phase III trials, leading to their FDA approval. However, their use has been associated with adverse events, including vascular inflammation and hemorrhage ^75^. A possible mechanism involves antibody binding to vascular Aβ deposits, triggering the complement cascade, which leads to membrane attack complex formation and subsequent blood vessel damage ^76^. Additionally, complement activation via C3a and C5a may stimulate microglia, further exacerbating vascular inflammation ^77, 78^. Both lecanemab and donanemab are humanized IgG1 antibodies, known for their potent effector functions that can trigger various immune responses and effector mechanisms ^79^. To enhance safety and efficacy, sdAbs represent a promising alternative. Their small size reduces immunogenicity and enhances tissue penetration, while their lack of an Fc region eliminates Fc-mediated effector functions such as antibody-dependent cellular cytotoxicity (ADCC) and complement-dependent cytotoxicity (CDC), potentially mitigating adverse effects ^80^. These properties position sdAbs as a safer and more effective approach for immunotherapy in neurodegenerative diseases.

We and others have demonstrated that antibodies can target proteinopathies both intra- and extracellularly ^1, 17, 22–25, 37, 81^. Antibodies capable of entering neurons can bind to protein aggregates within the endosomal-lysosomal and ubiquitin-proteasome systems, facilitating their clearance. Since tau pathology is predominantly intracellular, targeting tau within neurons is likely more effective than relying solely on extracellular clearance. However, neurodegeneration and aging can impair the function of lysosomes and proteasomes—the two primary intracellular degradation pathways ^6, 82–84^. Dysfunction in these systems may not only reduce the clearance of tau but also exacerbate toxicity by accelerating aggregate accumulation. These challenges underscore the need for strategies that enhance intracellular degradation pathways alongside targeted immunotherapies.

To address this, we developed an optimized sdAb, 1D9-LIRΔTP53INP2, designed to enhance tau clearance through the autophagy-lysosome pathway while maintaining high affinity for recombinant phosphorylated (p) tau and human-derived paired helical filament (PHF) enriched tau. This sdAb-based degrader was engineered based on the structure and function of Tumor Protein 53 Induced Nuclear Protein 2 (TP53INP2), which enhances recognition of LC3 on autophagic membranes and facilitates autophagy initiation. By integrating the LC3-interacting region of TP53INP2 with an anti-tau sdAb 1D9, we achieved efficient intracellular tau degradation. In both patient-derived neuronal models of tauopathy and tauopathy mouse models, the optimized sdAb demonstrated a substantial reduction in tau pathology. Notably, its therapeutic effects were accompanied by a significant decrease in astro- and microgliosis, indicating broader neuroinflammatory mitigation. These findings indicate that enhancing autophagic clearance via sdAb-based degraders represents a promising avenue for tauopathy treatment.

Following intravenous (i.v.) administration, we observed that 1D9-LIRΔTP53INP2 crossed the blood-brain barrier (BBB) efficiently and bound intracellularly to pathological tau. Notably, this sdAb successfully brought pathological tau into proximity with autophagosomes, facilitating the degradation of tau via the autophagy-lysosome pathway. Remarkably, six i.v. injections within six weeks led to a 27-60% reduction in brain p-tau levels and significantly improved motor function in tauopathy mice, further underscoring the therapeutic potential of this approach.

These findings establish our sdAb-based protein degrader as a compelling therapeutic strategy for clearing pathological tau in tauopathies. By coupling high target specificity with enhanced degradation mechanisms, this approach has the potential to address a critical challenge in neurodegenerative disease therapy. Future investigations will focus on evaluating long-term therapeutic efficacy and exploring its potential application to other tauopathies and neurodegenerative diseases.

To enhance tau degradation, two other approaches have been explored: fusing the RING domain of E3 ubiquitin ligase TRIM21 to either an anti-tau sdAb (Nanobody-RING) ^43^ or aggregation-prone tau (tau-RING) ^42^. Both strategies selectively degrade aggregated tau while sparing soluble tau. In tauopathy mouse models, adeno-associated virus (AAV)-mediated delivery of these constructs resulted in tau reduction and motor function improvement. Nanobody-RING may offer a safer approach due to its higher specificity. In contrast, tau-RING presents potential risks, as partial cleavage of the RING domain from mutant tau could generate pathological species. While AAV provides long-term expression and targeted delivery, its clinical translation remains challenging. The large human brain limits efficient distribution of therapeutic proteins, making widespread tangle clearance difficult. Although AAV delivery is more effective in smaller mouse brains, achieving sufficient transduction in humans poses a significant hurdle ^85, 86^. Moreover, AAV therapy is irreversible, raising concerns about long-term safety if adverse effects arise ^85–87^. Direct antibody injections present an alternative by enabling precise control over treatment, avoiding genetic modification, and reducing long-term risks. We recently demonstrated that three i.v. injections of a low-dose anti-α-synuclein sdAb or its proteolysis-targeting chimera (PROTAC) derivative (200 nmol/kg, ∼3 mg/kg) effectively cleared α-synuclein from the brains of synucleinopathy mice, with the PROTAC derivative showing greater efficacy ^88^. These findings highlight the potential of systemic antibody delivery for targeted protein clearance. However, the transient nature of antibody-based approaches may necessitate repeated antibody administrations, posing challenges for long-term treatment. Optimizing delivery strategies remains crucial for translating tau-targeted degradation therapies into clinical applications.

Additionally, both of the TRIM21 approaches discussed above utilized the ubiquitin-proteasome system (UPS) by tagging the catalytic RING domain of the ubiquitin ligase TRIM21 to facilitate tau degradation ^42, 43^. While these strategies yielded promising results, the intrinsic size constraints and substrate-conformation selectivity of the proteasome present significant limitations. Conformationally stable protein subspecies beyond the monomeric form are not only resistant to UPS-mediated degradation but may also obstruct proteasomal pores, leading to organelle overload and impaired degradation of other proteins ^44^. In contrast, autophagosomes offer a more robust degradation pathway, capable of processing oligomeric and aggregated forms of these pathological hallmark proteins ^44^.

This study primarily evaluated the therapeutic efficacy of the sdAb-based protein degrader in a patient-derived human neuronal cell model and a mouse model of tauopathy. Further studies are necessary to assess its safety, efficacy, and pharmacokinetic properties in more advanced preclinical systems. In addition, while no obvious toxicity was observed in our models, the long-term effects and potential risks of enhancing autophagosome-mediated tau degradation warrant further investigation.

### Conclusion

In conclusion, our study highlights the potential of sdAb-based protein degraders as a promising therapeutic strategy for tauopathy. By directly targeting tau for degradation, this approach offers a disease-modifying treatment with broad applicability to other neurodegenerative disorders driven by protein aggregation. Our findings support the feasibility of this precise and efficient degradation strategy, paving the way for future therapeutic development.

## Acknowledgments

We thank Dr. Peter Davies (Albert Einstein College of Medicine and Long Island Jewish Medical Center, Litwin-Zucker Research Center at Feinstein Institutes for Medical Research) for the tau antibodies PHF1, MC1, and CP27, which were obtained subsequently from Drs. Philippe Marambaud and Jeremy Koppel at the Feinstein Institutes. We are grateful to the NDRI (Philadelphia, PA), from which we purchased human brain tissue samples collected from patients with mixed AD/Progressive supranuclear palsy (PSP) pathology (Case ID: PSP-65484-01-slab 3). We are also grateful to the Tau Consortium iPSC line collection for providing the induced pluripotent stem cells (iPSCs) derived from fibroblasts of an individual carrying the autosomal dominant P301L mutation (Donor ID: F0510). We are grateful to Drs Kuo Min-Hao and Liu Mengyu from Michigan State for providing the recombinant hyperphosphorylated tau for the binding assay.

## Funding

This work was supported in part by National Institutes of Health (NIH) grants R21 AG058282, R01 AG032611, R01 NS077239, and R01 NS120488 (E.M.S.), and the Alzheimer’s Association Research Fellowships [AARF-22-924783 (Y.J.) and AARF-22-926735 (C.J.)].

## Author contributions

Y.J. performed most of the studies and related analyses. A. M. T. and Y.L. conducted the sectioning and immunohistochemistry staining of the mouse brains and related analyses. C.J. provided help with the iPSC culture study. K. F. L. and A. C. M. performed the Catwalk experiments. Y. L. maintained the animal colonies. E.E.C. generated the enriched PHF tau. R.P. and X.-P.K. helped with mammalian expression and purification of the sdAbs. Y.J. and E.M.S. designed the experiments and wrote the article. All authors had the opportunity to edit the article. E.M.S. supervised the project.

## Competing interests

E.M.S. is an inventor on a patent application related to this work assigned to New York University entitled Tau single domain antibodies, PCT/US2019/018571. Filed February 19, 2019. US Patent Application 20230203139 A1, Published June 29, 2023. The authors declare that they have no other competing interests.

## Data and materials availability

All data needed to evaluate the conclusions in the paper are present in the paper and/or the Supplementary Materials.

---

**Supplemental Figure 1:**
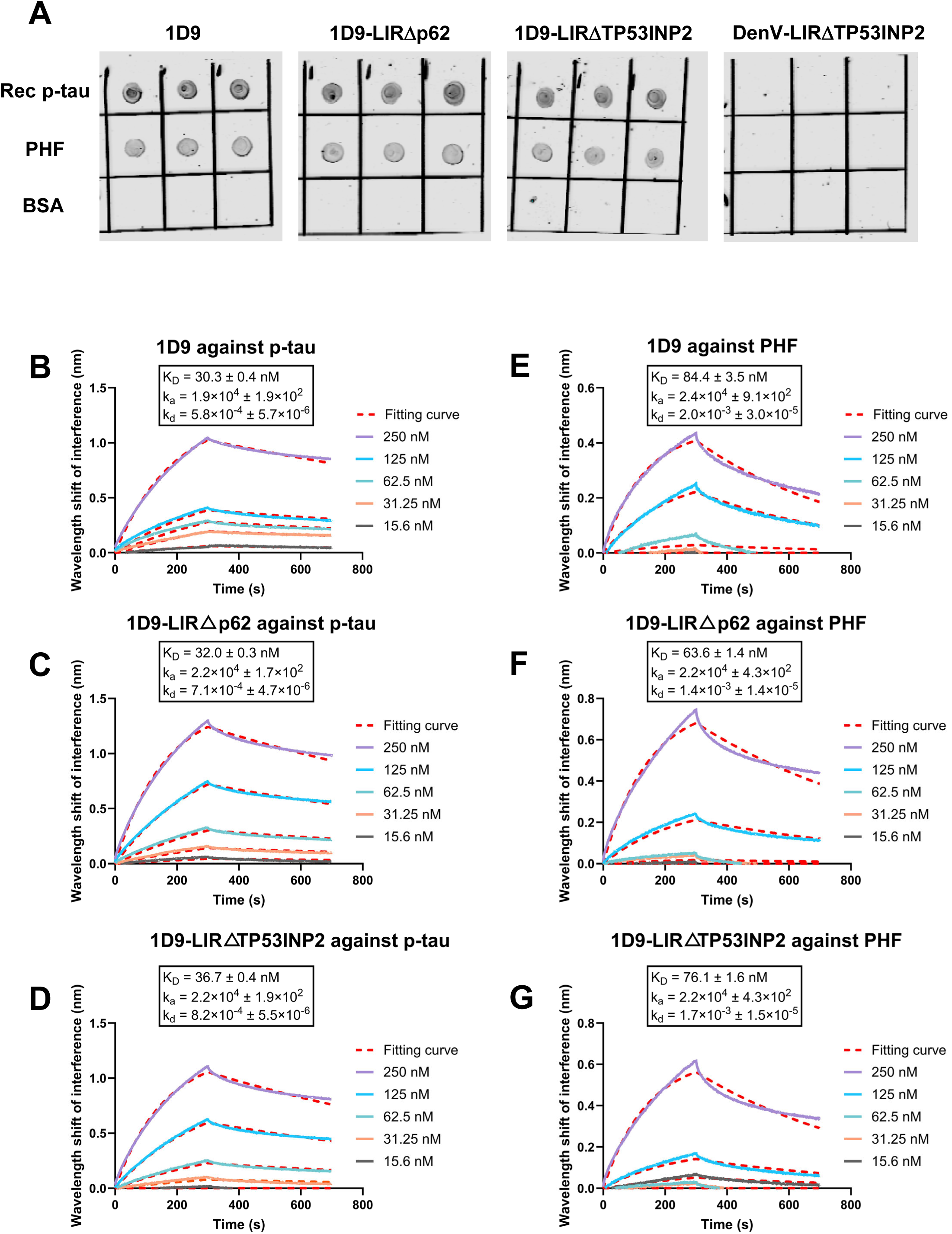
Binding analysis of single-domain antibody (sdAb)-based protein degraders to tau preparations. **(A)** Dot blot assay: Evaluation of tau binding capacity using recombinant phosphorylated tau (rec p-tau), patient-derived paired helical filament (PHF) enriched tau, and bovine serum albumin (BSA) as a control. **(B-G)** Binding kinetics analysis: Representative biolayer interferometry curves showing the wavelength shift (in nanometers, nm) for association and dissociation of sdAb constructs with tau preparations at various concentrations. The broken line represents the fitting curve used to calculate the dissociation constant (K_D_) ± standard deviation (SD) (*n* ≥ 3).

**Supplemental Figure 2:**
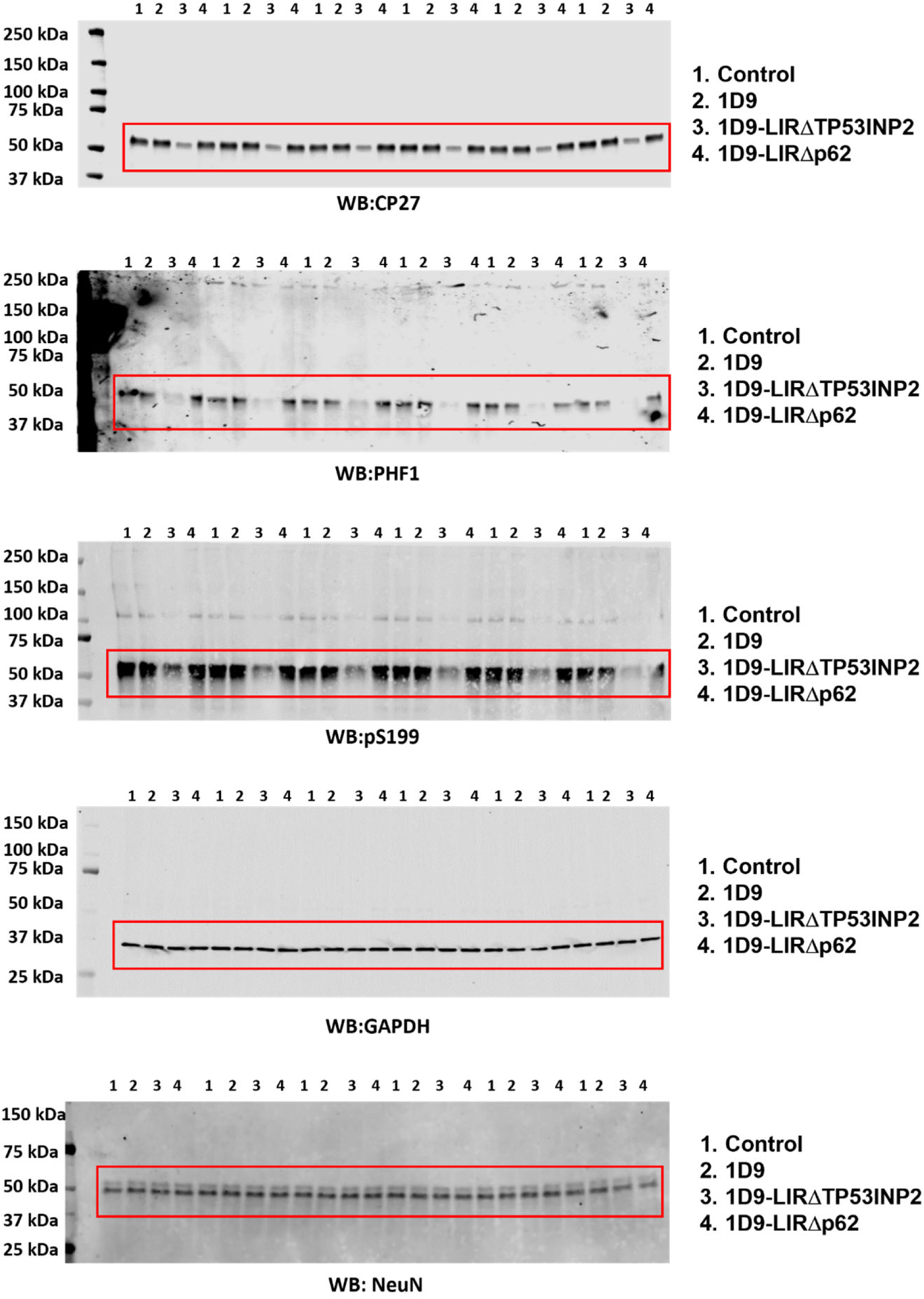
Complete western blots and bands quantified in Figure 2.

**Supplemental Figure 3:**
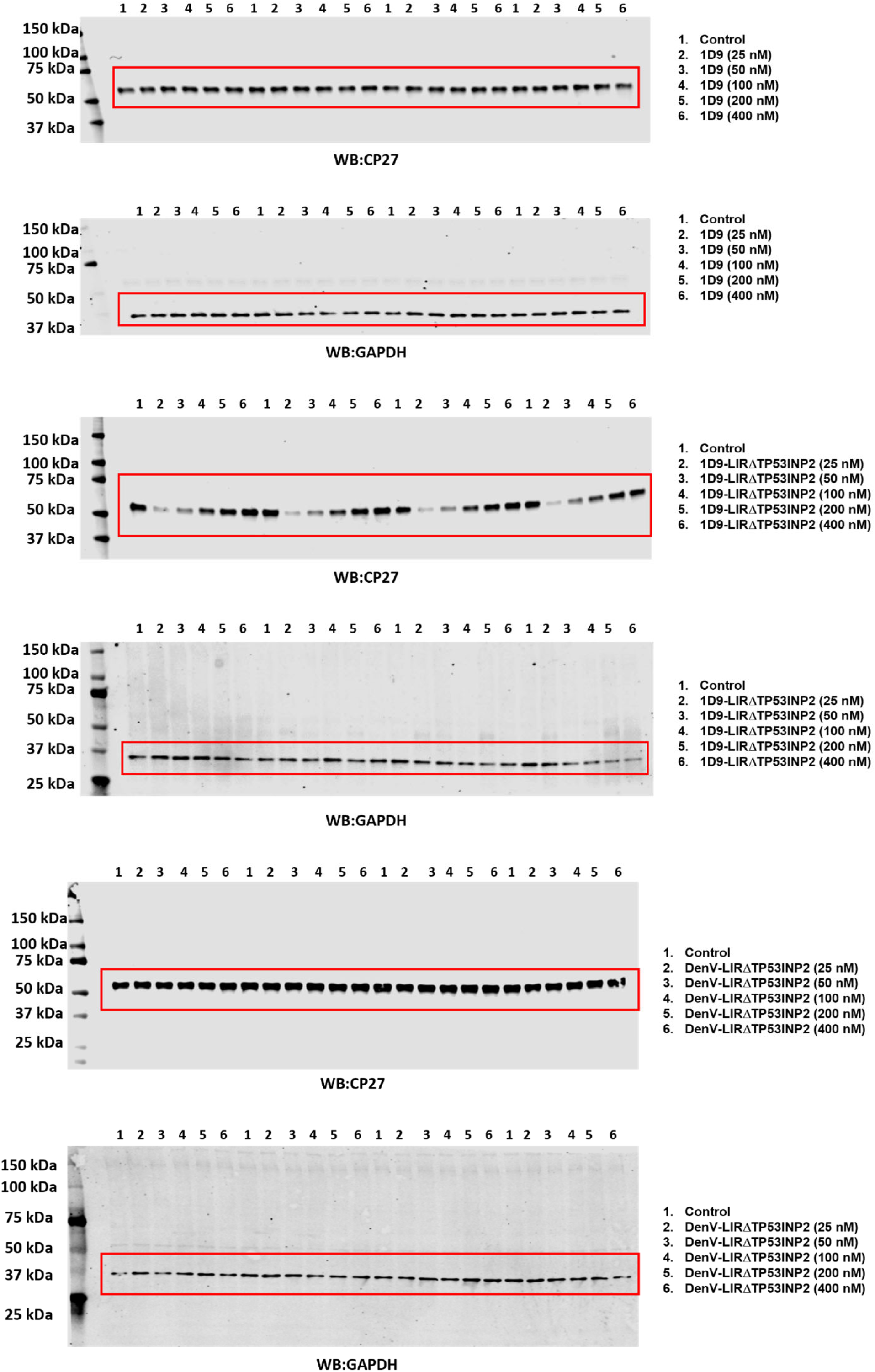
Complete western blots and bands quantified in Figure 3.

**Supplemental Figure 4:**
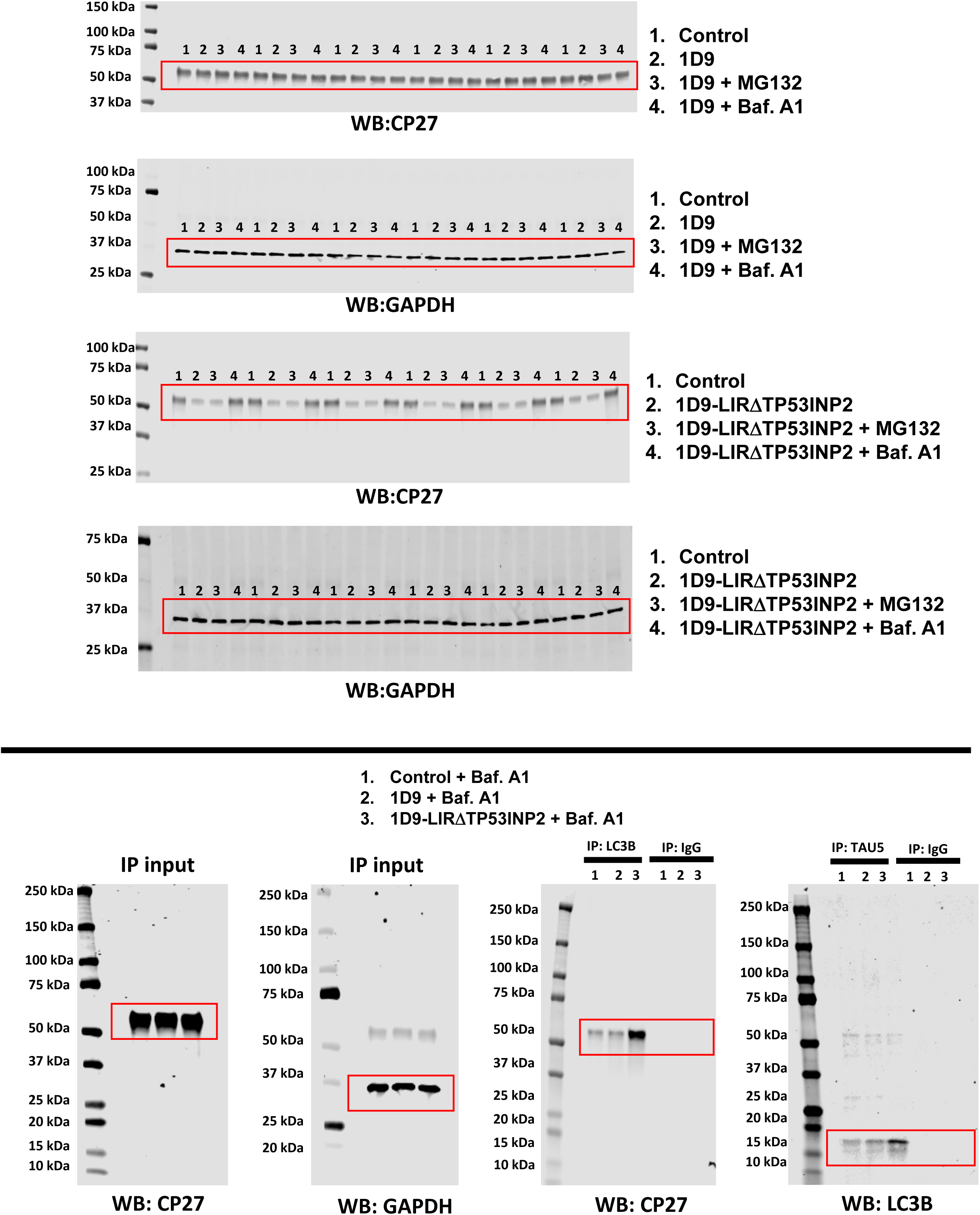
Complete western blots and bands quantified in Figure 4.

**Supplemental Figure 5.**
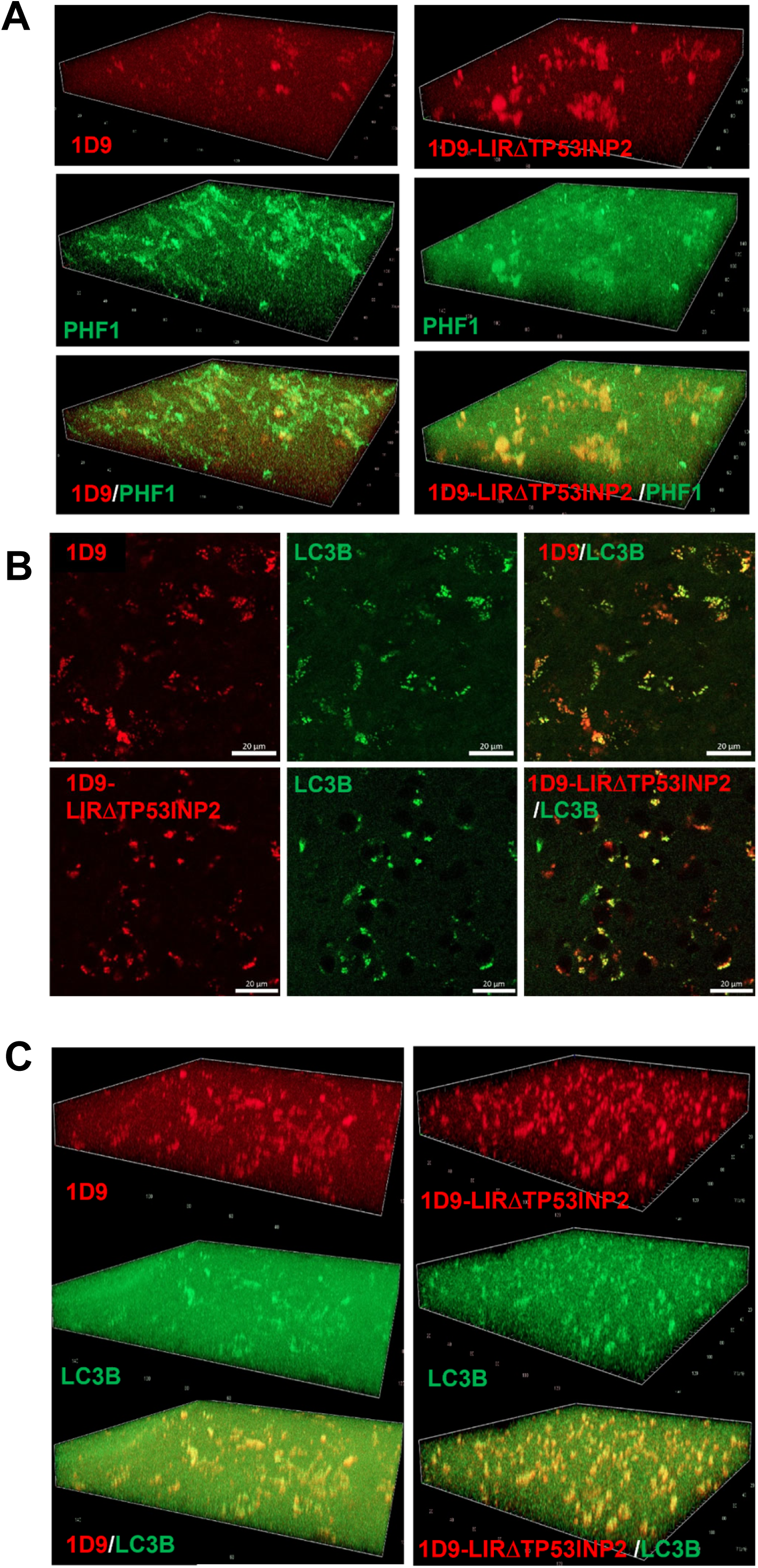
Colocalization of intravenously injected sdAb with phosphorylated tau (PHF1) and autophagosome marker (LC3B). **(A)** Z-stack confocal images demonstrate colocalization of sdAb 1D9 (left) and 1D9-LIRΔTP53INP2 (right) with phosphorylated tau (PHF1) in neurons. **(B)** Confocal images show colocalization of intravenously injected sdAb (1D9, top; 1D9-LIRΔTP53INP2, bottom) with autophagosome marker LC3B. Merged images reveal that both 1D9 and 1D9-LIRΔTP53INP2 entered the brain after an intravenous injection, were taken up into neurons, and colocalized with autophagosomes. Scale bars: 20 μm. **(C)** Z-stack confocal images confirm colocalization of sdAb 1D9 (left) and 1D9-LIRΔTP53INP2 (right) with LC3B.

**Supplemental Figure 6.**
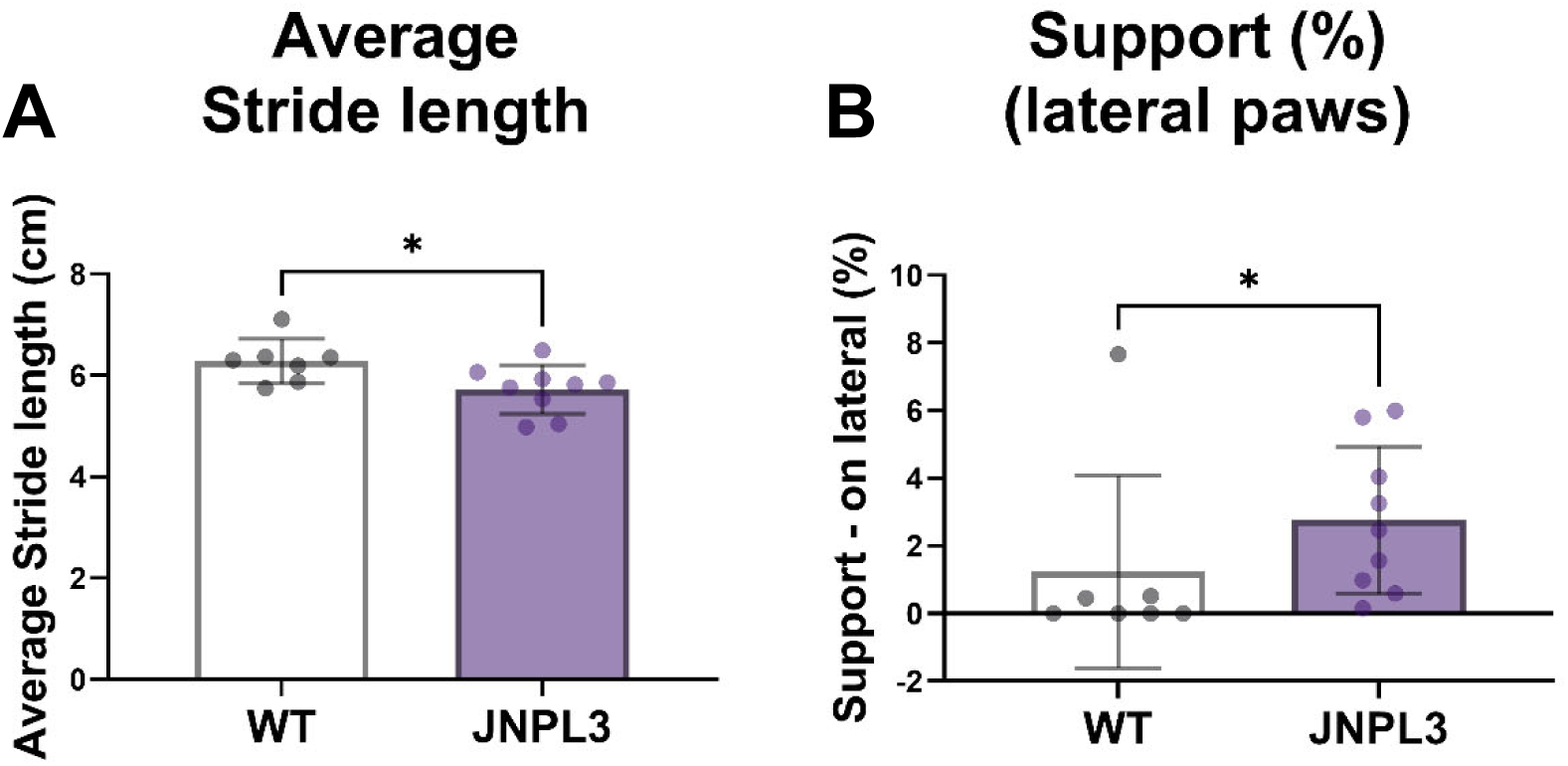
Gait abnormalities in JNPL3 female mice detected by CatWalk™ analysis. **(A)** Average stride length of all four paws was significantly decreased by 9.8% in JNPL3 female mice (*n* = 9) compared to age-matched wild-type littermates (*n* = 7), as assessed by CatWalk™ automated gait analysis (*F* [1,15] = 5.715, *p* = 0.0314). **(B)** The percentage of time spent on lateral support (defined as simultaneous contact of the ipsilateral fore- and hindlimbs) was significantly increased by 124% in JNPL3 mice (*n* = 9) compared to wild-type mice (*n* = 7) (*F* [3,23] = 3.301, *p* = 0.041).

**Supplemental Figure 7:**
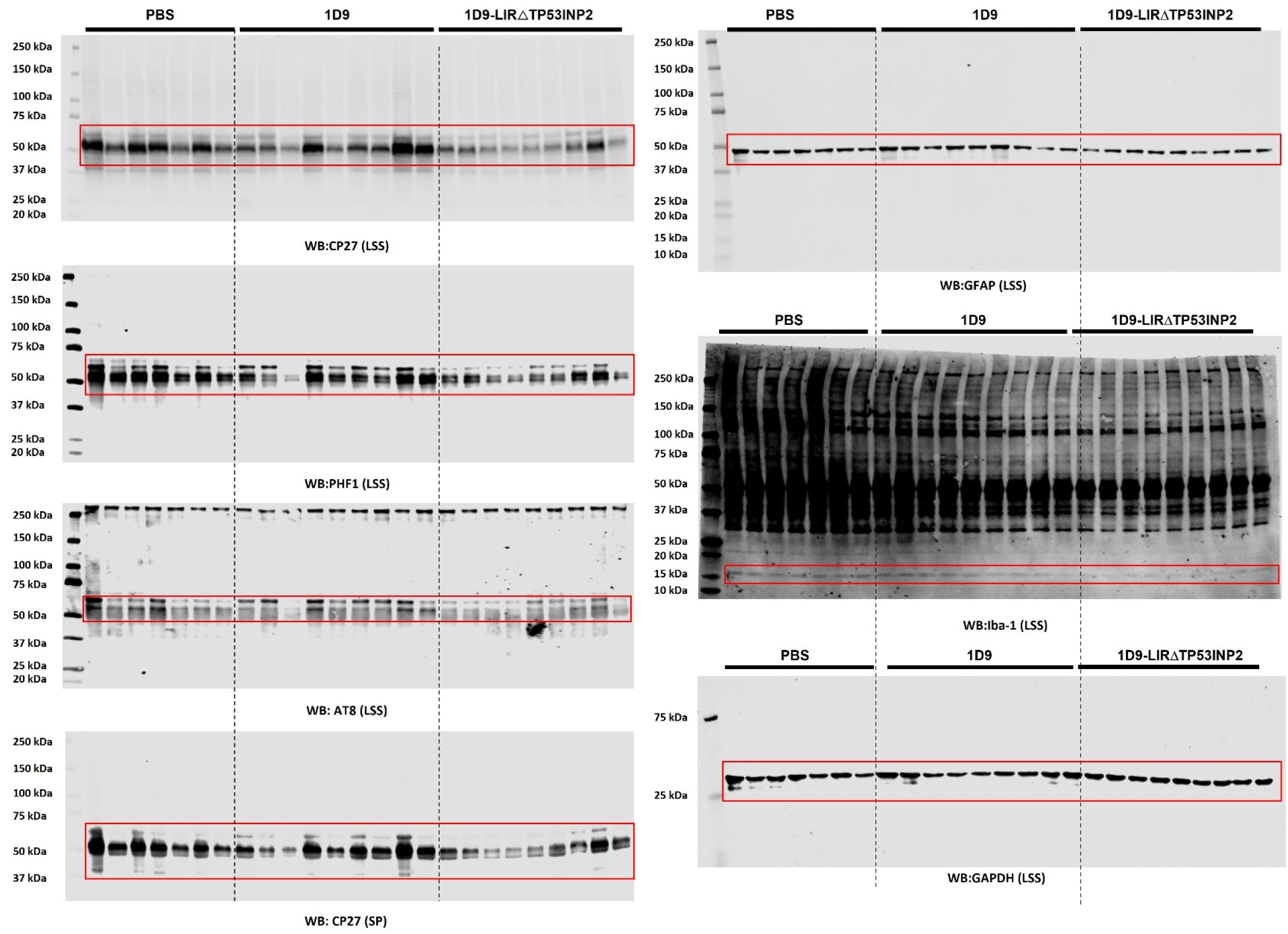
Complete western blots and bands quantified in Figure 6.

